# Indel variant analysis of short-read sequencing data with Scalpel

**DOI:** 10.1101/028050

**Authors:** Han Fang, Ewa A. Grabowska, Kanika Arora, Vladimir Vacic, Michael C. Zody, Ivan Iossifov, Jason A. O’Rawe, Yiyang Wu, Laura T Jimenez Barron, Julie Rosenbaum, Michael Ronemus, Yoon-ha Lee, Zihua Wang, Gholson J. Lyon, Michael Wigler, Michael C. Schatz, Giuseppe Narzisi

## Abstract

As the second most common type of variations in the human genome, insertions and deletions (indels) have been linked to many diseases, but indels of more than a few bases are still challenging to discover from short-read sequencing data. Scalpel (http://scalpel.sourceforge.net) is open-source software for reliable indel detection based on the micro-assembly technique. To date, it has been successfully used to discover mutations in novel candidate genes for autism, and is extensively used in other large-scale studies of human diseases. This protocol gives an overview of the algorithm and describes how to use Scalpel to perform highly accurate indel calling from whole genome and exome sequencing data. We provide detailed instructions for an exemplary family-based *de novo* study, but we also characterize the other two supported modes of operation for single sample and somatic analysis. Indel normalization, visualization, and annotation of the mutations are also illustrated. Using a standard server, indel discovery and characterization in the exonic regions of the example sequencing data can be finished in ~6 hours after read mapping.

## INTRODUCTION

Reductions in the cost of whole genome sequencing (WGS) and whole exome sequencing (WES), are opening the door for affordable sequencing of patients and the development of precision medicine [1]. Historically, genomic studies have focused on single nucleotide polymorphisms (SNPs) due to their high prevalence and relative simplicity to detect [12]. But recent advancements in sequencing technologies and computational methods have broadened the focus to include the role of insertion and deletion (indel) mutations. Indel mutations are defined by the addition or loss of one or more nucleotides of a DNA sequence. Frame-shift mutations are a highly disruptive class of indel mutations that alter the reading frame of protein coding sequences [2] and have been strongly implicated in neurodevelopment disorders, cardiovascular diseases, cancer, and many other human diseases [3-6]. In evolutionary analysis, the role of indels has been established and emphasized in both eukaryotic and prokaryotic genomes [7, 8]. In particular, studies have shown widespread occurrences of loss-of-function variants, especially indels, in protein-coding genes of human, plant and other species [9-11].

Recent studies have shown that indels are ubiquitous in human genomes, causing similar level of variation as SNPs in terms of total number of base pair changes, but with great diversity in size [13]. As the second largest group of variation in the human genome, there are typically more than one million small indels in the size range from 1 base pair (bp) to 100 bp per diploid genome compared to the human reference, with the majority of them being less than 10bp [14, 15]. The sizes of the indels in the human exome approximately follow a lognormal distribution with similar numbers of insertions and deletions [16]. However, indels are still very challenging to detect for multiple reasons: (i) long indels, especially long insertions are hard to detect with Illumina short reads, (ii) small scale repeats; short tandem repeats (STRs) and near-identical repeats increases the degree of ambiguity for mapping and assembly [17], (iii) non-uniform coverage distribution; irregularity in capture efficiency in exome sequencing and targeted re-sequencing can easily increase the number of false-negatives and false-positive calls depending on the type of study (e.g., de novo vs. single sample), (iv) sequencing and polymerase chain reaction (PCR) error; with PCR being especially error-prone around homopolymer A/T runs in the sequencing data [18] leading to the mapping/assembly problems described in (ii).

A common approach for variant calling (SNPs, indels, or otherwise) is to align reads one at a time to a reference genome, and to recognize when the reads disagree from the reference [19, 20]. Although this approach works well for SNPs, it is less reliable for indel detection. For example, reads supporting a long insertion will contain few bases matching the reference and will fail to map correctly. While reads supporting a deletion consist of bases from the reference, it may be hard to unambiguously map both sides of the deletion. In both cases the aligner may ignore parts of the reads (“soft-clip”) in order to place them on the reference or fail to map them at all.

Earlier methods for indel detection relied on paired-end and split-read information as a computational signature for the presence of an indel. Some tools such as GATK UnifiedGenotyper [19], SAMtools [21], and Dindel [20] use paired-end information to screen for indels where one read of a pair aligns well but the other pair does not. After identifying such regions, the algorithms use a local realignment of the reads to detect indels, although the sensitivity declines quickly for mutations longer than 5bp [18]. By using split-read information where the alignment for an individual read is split into two segments spanning structural variation breakpoints, methods like Pindel [22] and Splitread [23] are able to detect indels, especially deletions. Theoretically, this approach should be effective for deletions of any size, but the sensitivity is reduced due to the short read length of current sequencing technologies. Cortex, one of the first approaches for variant detection utilizing whole-genome *de novo* assembly with de Bruijn graphs, was reported to overcome such issues caused by short reads and alignment artifacts [24]. However, in practice this method is less sensitive than expected and accurate indel detection instead requires a fine-grained and localized analysis. Thus, in recent years, there has been much interest in developing specialized local assembly and microassembly methods [17].

One of the most sensitive and accurate approaches for indel detection from short read data is a micro-assembly algorithm, Scalpel. It was previously demonstrated to have substantially improved accuracy over eight algorithms including GATK-HaplotypeCaller [25] (v3.0) and SOAP-indel [26] (v2.01), while other methods report a large number of false negative calls [16]. In fact, Scalpel achieves very high accuracy (PPV=90%) of indel detection even on 30X WGS data (**Figure 1**). In this protocol, we describe the use of Scalpel for indel detection from whole genome and exome capture sequencing experiments. We introduce three different modes of indel detection: *de novo*, somatic, and single-sample for different study designs. First, the *de novo* mode is useful for calling germline *de novo* variants in nuclear families up to four people. Second, the somatic mode is useful for identifying somatic changes within matched samples, especially tumor/normal pairs in cancer studies. Finally, the single mode is useful for studies of a single proband.

**Figure 1.**
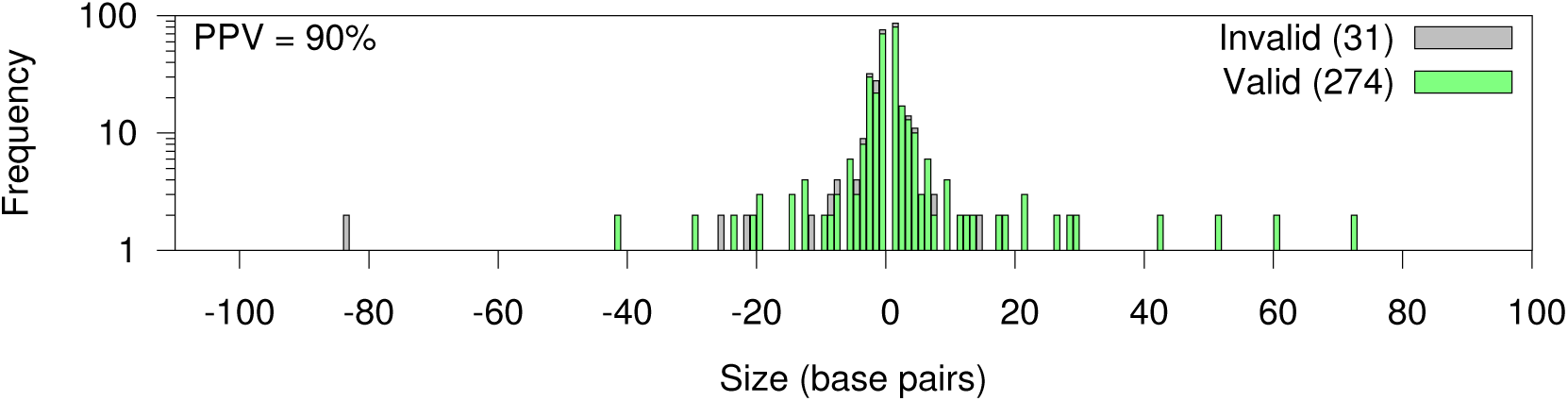
High accuracy of indel detection using Scalpel on WGS data. Scalpel was run in the single mode on 30X WGS data. This figure shows the size distribution of valid (green) and invalid (gray) indels that are randomly selected for validation (using targeted resequencing) in two previous studies. This validation set includes 160 and 145 candidate variants that were WGS-WES intersected and WGS-specific, respectively. Positive-predictive value (PPV) is computed by PPV=#TP/(#TP+#FP), where #TP is the number of true-positive calls and #FP is the number of false-positive calls.

### Overview of Scalpel micro-assembly strategy

Scalpel is a computational tool specifically designed to detect indels in next-generation sequencing (NGS) data. **Figure 2** outlines the main steps for the analysis of a sequencing dataset using Scalpel. To highlight the main focus of this protocol, the left panel of **Figure 2** depicts the specific scenario of detecting *de novo* indels in a quartet family composed of two parents and two children. We highly recommend reading the original Scalpel publication for a more extensive description of the method [16]. Here we briefly report the main ideas used in the micro-assembly strategy employed by Scalpel, and describe the new developments since the original publication of the software (v0.1.1 beta).

**Figure 2.**
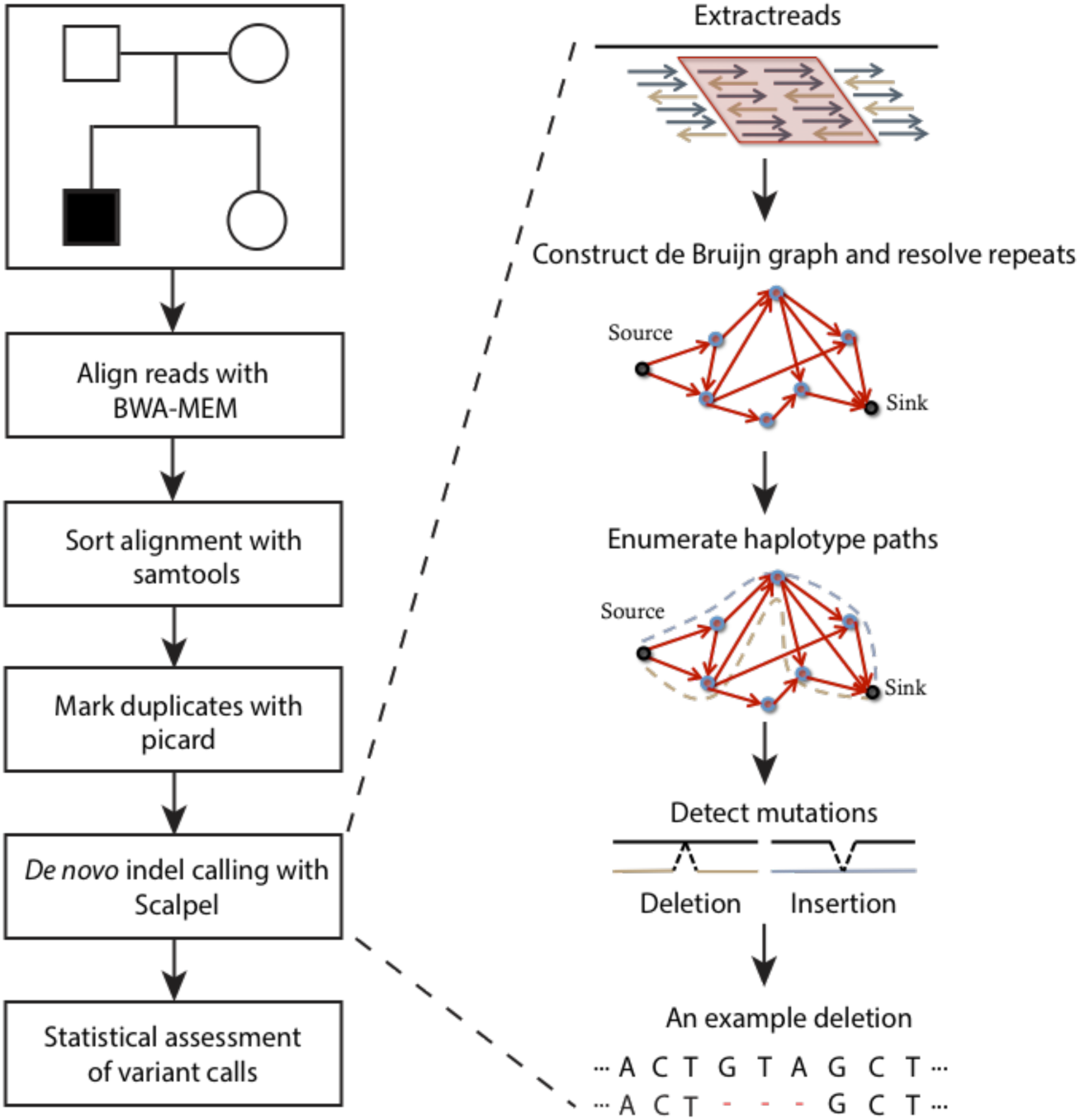
Main steps in the Scalpel protocol. Starting from raw sequencing data, reads are first aligned to the human genome using the BWA [34] software package. Following the standard practices in the field, the alignments are sorted (using Samtools [21]) and duplicates are marked (using Picard tools - http://picard.sourceforge.net). Note that, since Scalpel locally re-assembles the reads, this procedure is free of computationally expensive techniques such as indel realignment and base quality recalibrations. The BAM files obtained after the earlier steps are the input for Scalpel micro-assembly procedure.

Before running Scalpel, the sequencing reads (whole genome, whole exome, or custom capture) must be aligned to a reference genome using a short read mapping algorithm such as BWA-MEM, similar to the steps used for SNP calling or other analyses. It is worth noting that computationally expensive procedures like indel realignment and base quality recalibration are not necessary with Scalpel. Unlike in those analyses, the alignments are not directly used to find indels but instead are used to localize the analysis into computationally tractable regions. After alignment, Scalpel examines all the genomic regions provided in the input by the user in BED format (right panel, **Figure 2**). For each region, reads that align in the region or whose mates align in the region are extracted from the alignment and assembled independently of the reference using a de Bruijn assembly paradigm. If the size of a region is larger than the user-defined window size parameter, a sliding-window approach will be performed over this target region based on the window size and step size parameters. In order to reduce the number of errors in locally highly repetitive regions, Scalpel automatically performs a local repeat analysis coupled with a self-tuning *k*-mer strategy that iteratively increases the *k*-mer size until a “repeat-free” local assembly graph is built. In this context a repeat-free graph is a graph without exact repeats, which would introduce cycles in the de Bruijn graph, as well as near-identical repeats (up to 3 mismatches by default). The advantage of this strategy is that every genomic window will be analyzed using an optimal *k*-mer specifically tuned according to its sequence composition. The graph is then exhaustively explored to identify end-to-end paths spanning the selected region. These paths, representing *de novo* assembled sequences of the short reads, are then aligned to the reference window to detect candidate mutations using a sensitive gapped sequence aligner based on the Smith–Waterman algorithm.

Scalpel supports three modes of operation: *single, de novo,* and *somatic.* In the single mode, Scalpel detects indels in one single dataset (e.g., one individual exome). In the *de novo* mode, Scalpel detects *de novo* indels in a quad family (father, mother, affected child, unaffected sibling). In the somatic mode, Scalpel detects somatic indels from the sequencing data coming from matched tumor and normal samples. In this protocol we illustrate the use of Scalpel by focusing on the *de novo* mode; however we describe the alternative and advanced operation modes in **Box 2** and **Box 4**, including a discussion of the computational requirements for running Scalpel and how those differ between whole-genome and whole-exome studies. **Box 3** provides further guidelines on how to export and filter the mutations based on coverage and quality scores and how those operations could impact sensitivity and specificity.

##### Box 1: Representing indels

Unlike SNPs that are always represented with a unique genomic coordinate and base substitution, an indel can have an ambiguous representation. For example, if there is a 1 bp deletion in a long homopolymer (…AAAAAA…), deleting any A will give rise to the same haplotype but just with a different position. A more complex example which gives rise to two logically equivalent 3 bp deletions is shown in **Figure 8**:

**Figure 8.**
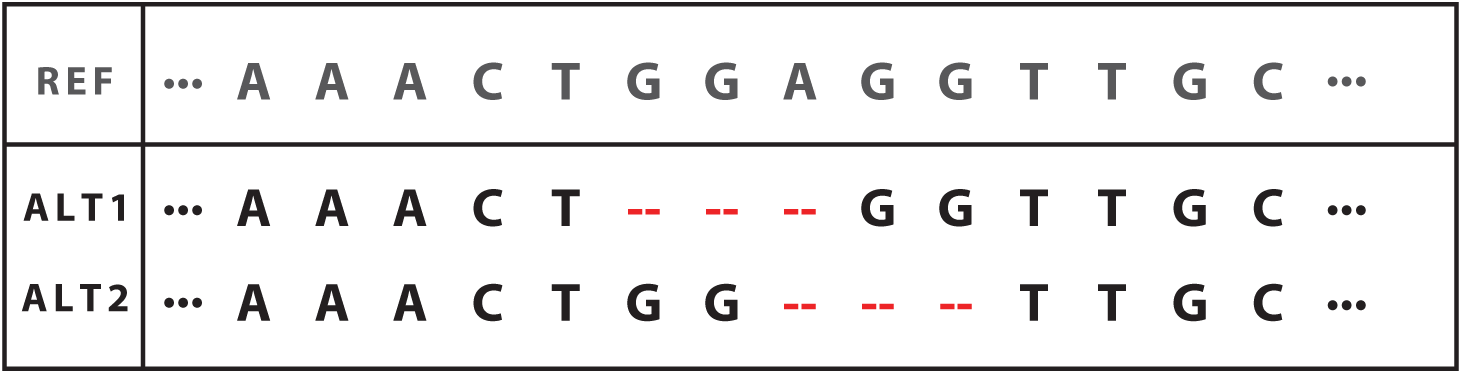
An example indel representation ambiguity due to normalization. Alt1 represents a left-normalized indel (employed by Scalpel) while Alt2 represents a right-normalized indel.

Note that two different 3bp sequences can be deleted (GGA or AGG) at two different locations generating the same alternative sequence. The solution to this ambiguity is to consistently *left-* or *right*-normalize the signature of the mutations. This operation consists of shifting the start position of the mutation to the left (or right) as long as the resulting sequence (after the deletion or insertion of the specific number of bases) is still the same as the one generated by the original mutation. Note that at the end of this process the new signature for the indel can have a new coordinate as well as a new (deleted or inserted) sequence but the size must remain the same as the original. For example in the case of the previous 3bp indel, the deletion is shifted to the left by 2 positions and a new 3bp sequence (GGA) is deleted.

Since different methods might report different signatures for the same indel, it is essential to consistently normalize the signature (typically left-normalization) when comparing indels called by different tools or querying different databases (dbSNP, 1000G, OMIM, etc.). Scalpel always returns a list of variants that are left-normalized. However, if it is unclear what representation has been used for a set of variants made by other callers, there are different tools available that can normalize a list of input variants, including *vt normalize* [37], *bcftools norm*^1^, *GATK LeftAlignlndels*^2^. Indel normalization is now becoming standard practice and widely used variant annotation software, such as ANNOVAR [38], is now enforcing left-normalization as the default representation for indels. Updated variants databases (e.g., 1000 Genome Project, dbSNP, etc) were made available and ANNOVAR users are highly encouraged to left-normalize (using any of the previously listed tools) the variants prior to annotation. However, note that some of the databases use right-normalization or normalize relative to the sense of the transcript, so users are encouraged to refer to the documentation of each tool separately.

##### Box 2: Alternative operation modes and computational requirements

Scalpel is designed for UNIX-type operating systems, and provides a command-line interface. Users are expected to have a basic familiarity with operating in a UNIX environment. The discovery pipeline of Scalpel is executed from the command line via a master (perl) script called “scalpel-discovery” and it requires a minimum number of parameters describing the input alignment files (BAM format), the reference genome (FastA format), and the target region (BED format) to analyze.

Scalpel supports three operation modes: *single, somatic,* and *denovo.* The denovo mode is described and used in the procedure of this protocol. Here we report the basic usage and command lines parameters for the other two operation modes. To call variants on one single sample (e.g., single exome or single whole-genome dataset), use the following command:

~~~
$ scalpel-discovery --single --bam *file.bam* --bed *regions.bed* --ref *genome. fa*
~~~

where *file.bam* is the bwa aligned BAM file of the reads (after sorting, indexing and PCR duplicates marking), *regions.bed* contains the set of target regions in BED format (typically the list of exonic coding regions), and *genome.fa* is the reference sequence in FastA format. It is important to provide to Scalpel the same reference file that was used to align the reads in the BAM file.

If calling variants on a tumor/ matched normal pair, execute Scalpel as follows:

~~~
$ scalpel-discovery --somatic --normal *normal.bam* --tumor *tumor.bam* --bed *regions.bed* --ref *genome.fa* --two-pass
~~~

where *regions.bed* and *genome.fa* are the same files as described before; *normal.bam* and *tumor.bam* are the bwa aligned BAM files of the reads for the normal tissue and the tumor tissue sample respectively. Also note the use of the *“--two-pass”* option which enables Scalpel to perform a second round of indel verification on the candidate list of somatic mutations to reduce the number of false-positive calls. For example, in the case of a tumor/normal pair, a more sensitive analysis is performed on the normal sample to identify any signature of the candidate mutation in the tumor that was missed during the first pass of the analysis. We highly recommend using the *“--two-pass”* option for *de novo* and somatic studies. Exceptions to this rule are studies with extremely high coverage (e.g., 1000x or more) that can be obtained for example in panel studies of cancer samples, for which the use of the two-pass option is not required.

It is best to run Scalpel on a multicore computer with at least 64GB of RAM. The relative computational requirements depend on the type of data to analyze. For example, in the case of whole-exome analysis, 10 CPUs and a minimum of 10Gb or RAM will be enough to perform the analysis in a few hours. In the case of whole-genome study instead, in order to reduce its memory requirements, it is recommended to run Scalpel on each chromosome separately and then merge the lists of detected call sets. Given the more uniform coverage distribution of whole-genome data and the increasing read length of Illumina technology, we also recommend to increase the window size (default 400 bp) to 600 bp or larger. For example, the following command can be used to call variants on chromosome 22 using 10 CPUs:

~~~
$ scalpel-discovery --single --bam *file.bam* --ref *genome.fa* --bed 22:1-51304566 --window 600 --numprocs 10
~~~

##### Box 3: Exporting variants and filtering considerations

By default Scalpel exports the list of detected indels in a VCF file within the selected output directory according to the parameters. However it is recommended to explore different filtering criteria using the export tool (scalpel-export). The raw list of mutations detected by Scalpel is always available in a database within the output directory that can be queried with the export tool using the following command:

~~~
$ scalpel-export [single|somatic|denovo] --db *database.db* --bed *regions.bed* —ref *genome.fa* [options] > *variants.vcf*
~~~

All detected mutations are exported, but following the standard practices of the VCF format, high quality mutations are flagged as “PASS” in the FILTER column. For non-PASS mutations the FILTER field contains the list of filters that were applied to the variant explaining the reason why the variant was considered of low quality. For example, in the VCF snippet below, the first indel is of high quality and is labeled as “PASS" while the second indel does not satisfy the minimum phred-scaled Fisher’s exact test score requirement and it is flagged as “LowFisherScore”:

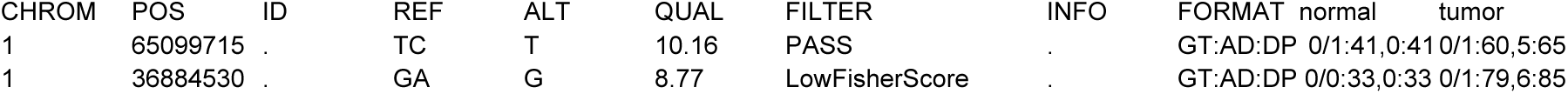

These filters can be further controlled using some of the command line parameters (**Table 2**). Similarly to the scalpel-discovery command, the export tool requires the mode of operation to be specified according to type of study (single, somatic, denovo). Different parameters and filters are available for each operation mode; here we discuss some of the most important ones and give recommendations on how to adjust them according to the type of study.

*VAF and support.* High number of supporting reads and high VAF are typical signals of strong evidence for a variant. All three modes of operation provide parameters to control the thresholds used for the minimum number of supporting k-mers *(--min-alt-count, -- min-alt-count-tumor, --max-alt-count-normal)* and minimum allele fraction for the alternative allele *(--min-vaf, --min-vaf-tumor, --min-vaf-affected)* used to filter low quality variants. Although default values are provided, optimal cutoffs for these numbers depend on the coverage available for the sample and the type of study (single, somatic, denovo). For example, since somatic calls are typically found at lower VAFs in the data, the user may need to adjust these parameters according to the level of purity, ploidy and clonality (if available) of the data.

*Contamination in normal samples.* The matched normal sample to the tumor is typically assumed not to be contaminated with the tumor sample. However, in practice, it is possible to have a very low level of contamination of the tumor in the normal sample as well. In this scenario two parameters, *“--min-alt-count-tumor"* and *“--min-vaf-tumor”,* (which by default assume no contamination in the normal) can be used to allow mutations in the tumor, which are also found in the normal at low allele fraction, to be called as somatic.

*Statistical test and scoring.* For germline (inherited and de novo) mutations the relative balance between the alternative and reference counts is estimated using the chi-squared test statistic. The cutoff used can be adjusted via the *“max-chi2-score”* parameter. A larger value will increase sensitivity but produce a larger number of false-positives. We recommend using chi-squared score < 20 to export high confidence indels. Differently from germline mutations, which are expected to be relatively balanced in their reference and alternative counts, somatic variants are usually out of balance due to several known problems with cancer data (e.g., ploidy, clonality, purity). The Fisher’s exact test is generally used to determine if there are nonrandom associations between the allele balances in the tumor and the normal. Scalpel internally scores the mutations by computing a phred-scaled p-value Fisher’s exact test score and the filtering cutoff can be adjusted via the *“--min-phred-fishef’* parameter. By default this parameter is set to 10, but lower values will increase sensitivity at the cost of specificity.

**Table 1.** Comparisons and validation rates of indel detection with WGS and WES.

|  | Indels | Valid | PPV | Indels<br>(>5 bp) | Valid<br>(>5 bp) | PPV<br>(>5 bp) |
| --- | --- | --- | --- | --- | --- | --- |
| WGS-WES<br>intersection | 160 | 152 | 95.0% | 18 | 18 | 100% |
| WGS-specific | 145 | 122 | 84.1% | 33 | 25 | 75.8% |
| WES-specific | 161 | 91 | 56.5% | 1 | 1 | 100% |

Note: The validation rate, positive predictive value (PPV), is computed by the following: $PPV = \#TP / (\#TP + \#FP)$ , where #TP is the number of true-positive calls and #FP is the number of false-positive calls.

**Table 2.**
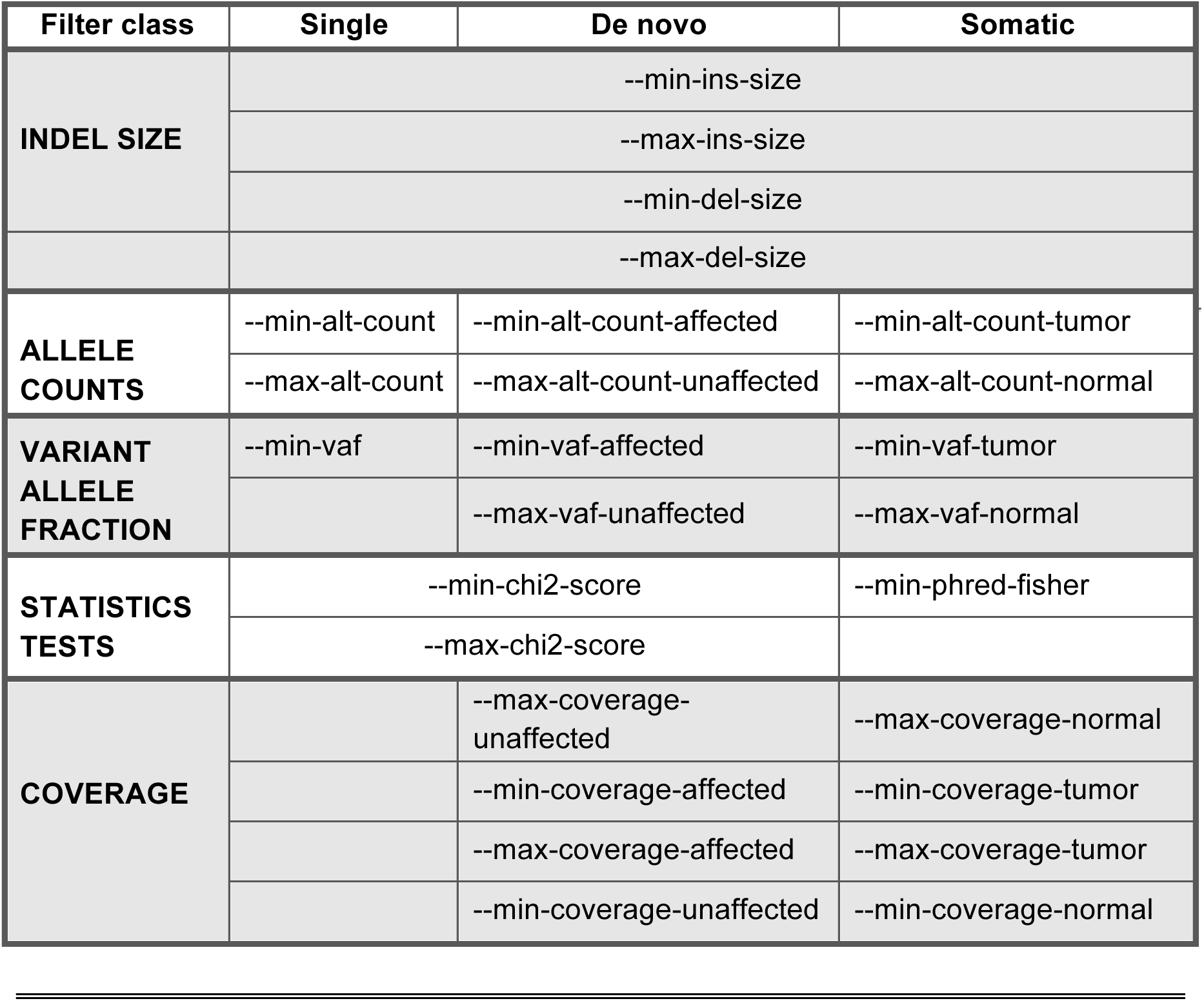
List of available parameters for each operation mode used to filter low quality variants (scalpel-export).

##### Box 4: Advanced Scalpel operations

Variant calling is a computational step widely applied to a multitude of different projects and datasets, and, as expected, the default parameters of the tool cannot handle all situations equally well. Here we describe how to adjust the optional parameters accordingly in a few common scenarios.

*Deep sequencing.*. Some projects require deep sequencing to allow, for example in cancer, the detection of low allele fraction mutations. With higher coverage there is also enrichment of the errors in the data. In the most extreme scenarios of very deep sequencing experiment with thousand fold coverage or greater (e.g., cancer gene panels), these errors contribute to an increased complexity of the assembly graph to the point where the associated region to assemble would be discarded. Thus, it become necessary to increase the minimum *k*-mer coverage used to remove low-coverage nodes (parameters *“--lowcov”* and *“--covratio”).* This can typically solve the problem by reducing the complexity to a level that the graph can be efficiently analyzed. Also, by default regions that have coverage >10,000 are not processed. If higher coverage is expected, the maximum average coverage allowed per region must be adjusted accordingly *(“--maxregcov”* parameter).

*Detecting very long indels (>100bp).* There are cases where the researchers might have some evidence or prior knowledge about the presence of larger indels (greater than 100 bp and up to 1 kb) in their data set. In this scenario Scalpel can be used to genotype these loci for the presence of the mutations. Two parameters can be adjusted to handle and improve the sensitivity in such cases:

– *“--coords”:* using this parameter the user can specify a list of selected coordinates to examine. The expected format is a tab-delimited list of chromosome names and positions.
– *“--window”:* by default the list of positions are analyzed using a window size of 400 bp. For indels approaching the window size or larger, it is necessary to increase this parameter to allow for enough unique sequence on both sides of the mutation. For example in the case of a 400bp insertion it is recommended to use a window size of at least 600 bp so that 100 bp of unique sequence can be used to anchor the mutation on the reference on both sides.

*Inspecting the assembly.*

In same special cases the user may be interested in examining the final assembly generated by Scalpel. This information is stored internally by the program in the log files. By default the log files are not saved in the output directory since they can be significantly large in size, especially for whole genome analysis. It is possible, however, to change this behavior by using the *“--logs”* option, but we recommend doing so only if a relatively short list of small regions are being analyzed. The file contains detailed information describing the different stages of the assembly. It is out of the scope of this paper to describe the complete format of the log files; instead we will focus on the section containing the final assembly and alignment of the region of interest. A typical alignment of the assembled sequence to the reference will look like the following one:

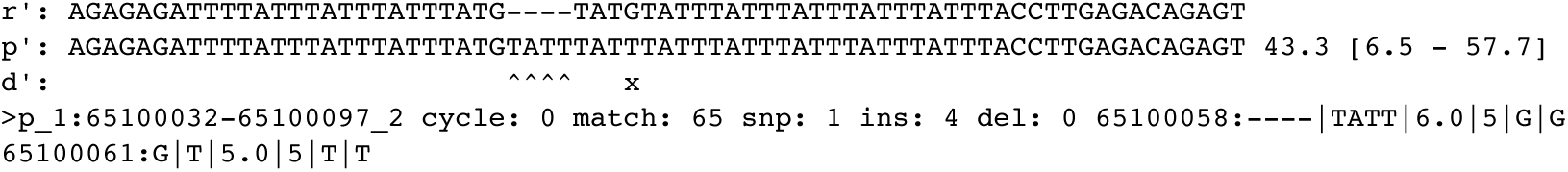

where `r’` is the reference sequence, `p’` is the sequence assembled by Scalpel followed with information about the minimum and maximum coverage across the assembly, and `d’` is the alignment string showing the differences between the reference and the assembled sequence. In this case the assembly contains a complex mutation composed of an insertion of 4 bases (`TATT`) together with single base substitution (`G->T`). The last line reports the genomic coordinates of the region followed by: 1) number of cycles detected `(cycle: 0), 2)` total number of matches to the reference (`match: 65`), 3) number of SNPs, insertions and deletions (`snp: 1 ins: 4 del: 0`), and 4) a list of signatures describing each mutation. The signature starts with the position of the mutation followed by “:” and a list of fields separated by the symbol “|” (e.g., 65100058:028050|`tatt|6.0|5|G|G`). In order each field contains:

– Position
– Reference sequence
– Alternative sequence
– Average coverage supporting the mutation
– Minimum coverage supporting the reference
– Base pair preceding the mutation in the reference sequence
– Base pair preceding the mutation in the alternative sequence

In the first version of scalpel (v0.1.1), all possible paths in the final graph were exhaustively examined using a breadth-first-search traversal approach. This strategy worked well for the majority of the human genome with limited numbers of mutations leading to the generation of one or two paths. However this step is computationally expensive for a small number of regions with high level of heterozygosity or higher sequencing error rate that generate exponentially many alternative paths due variants not linked by the same k-mer. Since the release of a new version (0.4.1), Scalpel instead enumerates only the minimum number of source-to-sink paths that cover every edge of the graph using a network flow approach. This strategy still detects all the mutations in the graph but significantly reduces the computational requirements by aligning to the reference a much smaller set of paths. Another important addition in the new version of Scalpel is the ability to better handle regions characterized by sudden drops in coverage. After removal of low-coverage nodes, the de Bruijn graphs associated to these regions can be disconnected into multiple connected components, which are now analyzed independently.

Other software packages also implement similar localized sequence assembly strategies to the one employed by Scalpel. These include GATK HaplotypeCaller [25], SOAPindel [26], Platypus [27], ABRA [28], TIGRA [29], DISCOVAR [30], and Bubbleparse [31]. Although they all employ a local read assembly step, these tools differ in how they explore the graphs and in their relative ability to handle repeat structures. Further, although some tools can perform multi-sample calling, none of these tools provide direct support for *de novo* or somatic mutation calling. We also encourage the users to read a review on the challenge of small-scale repeats for indel discovery for a more in depth discussion of the differences [17].

### Limitations of the protocol and software

Scalpel provides several advantages to standard mapping approaches but, like any bioinformatics algorithm, it does not attempt to address all possible types or sizes of mutations at once. In our experiments Scalpel was able to reliably detect deletions up to 400 bp (including deletions of *Alu* mobile elements) and insertions shorter than 200 bp, but the sensitivity is reduced for longer indels given the available read lengths (data not shown). Even within this size range, Scalpel, and all pipelines, has lower sensitivity for indels in low coverage regions that are supported by very few reads. In the worst scenario, a combination of low coverage within a complex repeat region may require a *k*-mer size too large for assembling across the mutation, introducing false negatives. Phasing of the discovered mutations is not supported and, given the locality of the assembly, it would be possible to phase only mutations within the same window (400bp by default). Thanks to the new advances in long-molecule sequencing technologies (e.g., 10X Genomics), in the near future it will be possible to combine such technologies for phasing mutations hundreds of kilobases to megabases apart. For variant calling purposes, it is ideal to have a high quality reference genome available. This is also true for indel calling with Scalpel because assembly errors might greatly increase the number of variants and the read localization will not be effective unless a complete representation of the genome is available. Users working with data from a genome without a reference should first generate a high quality assembly using one of the several whole-genome assemblers [32, 33]. This procedure can be easily adapted to work with a draft assembly, but no testing has been performed and the results could be unpredictable. Tumor/normal and multiple family members can be analyzed together, but joint calling across a large number of samples is not support by Scalpel. This protocol also assumes that sequencing was performed using the Illumina sequencing platform, including MiSeq, HiSeq 2000, and HiSeq X sequencers. Other sequencing technologies (e.g, Ion Torrent, Sanger, SOLiD) can be also used for studies like the one reported here, but the software pipeline used in this protocol does not support them. No graphical user interface is available for the steps performed in this protocol; all the operations are performed through the UNIX shell. Some of the tools used here, such as BWA and Picard tools, are now available through cloud-based web interface systems such as Galaxy (https://usegalaxy.org/). We look forward to seeing Scalpel integrated into such systems in the near future.

### Overview of the protocol

In the Materials section, we present a step-by-step protocol for identifying *de novo* variants in a HapMap family from PCR-free Illumina HiSeq2000 data. Here, we provide an overview of using Scalpel to discover *de novo* and inherited indel mutations within a quad family of two parents and two children, one affected and one unaffected with a certain phenotype. But internally within the algorithm the two children are treated identically, which can support additional use cases. The input to the algorithm can be data from WGS, WES, or targeted sequencing experiments. A two-pass search mode is employed by Scalpel when calling *de novo* or somatic mutations. In the first pass, Scalpel identifies indels in each of the samples using parameters designed to balance between sensitivity and specificity. In the second pass, Scalpel performs a more sensitive search in the parents for the indels identified in the children to reduce false positives *de novo* calls in regions of low coverage in parents. We also show how to extract indel calls that fall into target regions and filter out false-positive calls with respect to their sequence composition and variant quality (**Figure 3**). Finally, we present one of the available methods for annotating the mutations, to identify any potential disease-related mutations. The protocol provides general guidelines for standard operations required to analyze and evaluate indel calls. We also illustrate several sources of indel calling errors, which could be introduced by library construction, sequencing or alignment. Whenever possible, visualization of data/results is performed using IGV alignments, and auxiliary scripts are provided for plotting size, allele fraction distribution, etc.

**Figure 3.**
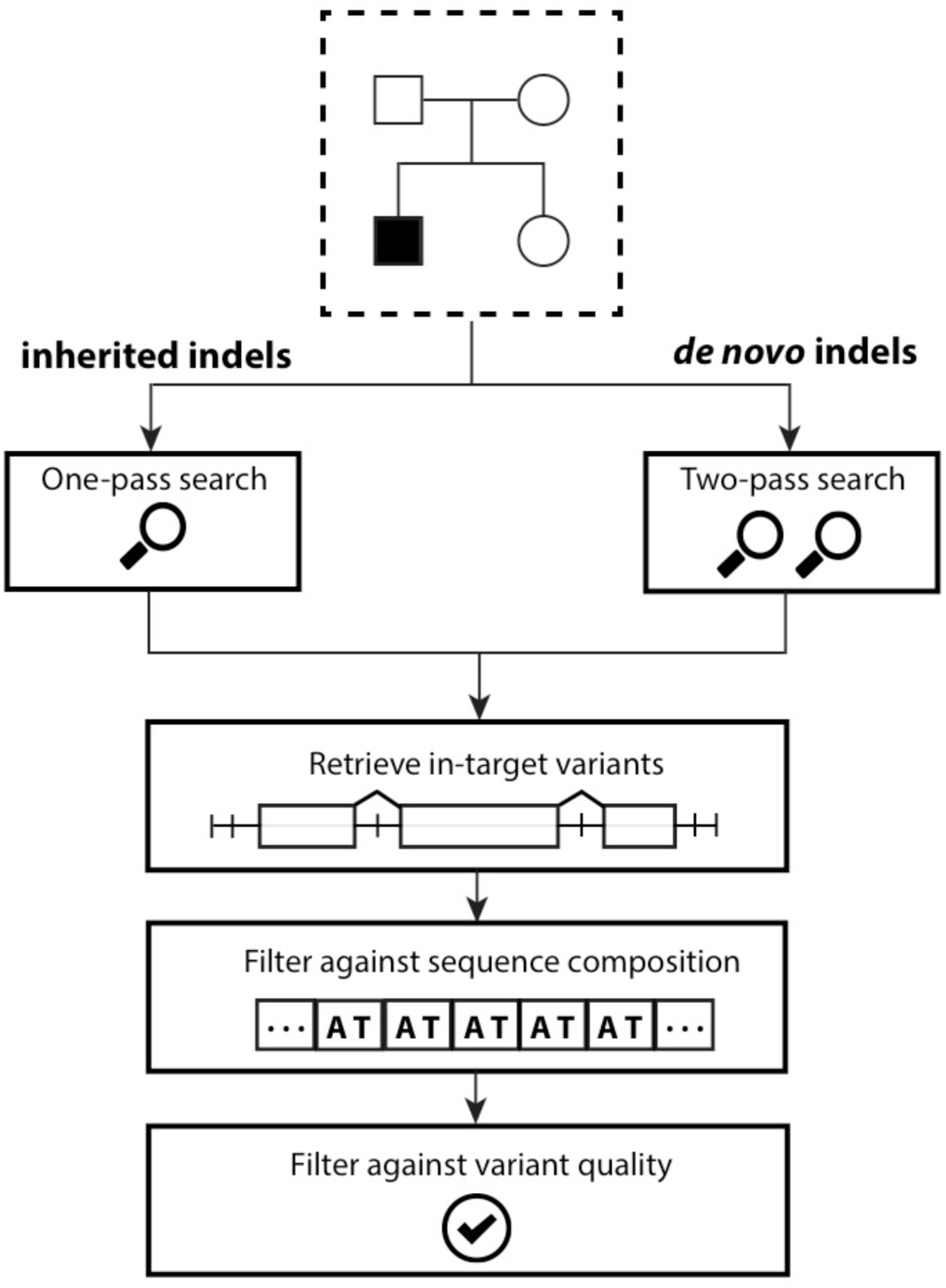
Overview of the variant filtering cascade. This figure shows the filtering cascade used to report high quality *de novo* and inherited indels within the target region; coding regions in this case. (1) Inherited and de novo indels are analyzed separately; (2) only variants within the target regions are exported; (3) Low quality indels are identified and removed based on sequence composition (e.g., STRs); (4) Additional filters based on supporting coverage and allele balance are used to reduce the number of false-positives.

This protocol is based on the use of version v0.5.1 of the Scalpel software. Users should keep in mind that the software is continuously under development and some of the parameters, file names, output formats could change in the new releases of the software. The most recent version of the code and documentation is always available at http://scalpel.sourceforge.net. This protocol follows very closely a typical usage of the software, however we recommend that the users perform the full procedure described herein before running the pipeline on their own data.

### Experimental Design

In this protocol we use publicly available WGS data to detect and analyze indels within a family. However, when designing a new study, researchers are typically faced with the problem of choosing suitable sequencing and bioinformatics strategies to answer the relevant scientific questions. There are many factors that play a role in study design, including depth of coverage, read length, parameter tuning, WGS versus WES protocols, the use of PCR amplification, cost per basepair, etc. In this section, our goal is to provide some guidelines on the impact of such different experimental design choices on the sensitivity and accuracy of indel detection.

### Whole-genome vs whole-exome sequencing

Although WES is a cost-effective approach to identify genetic mutations within the coding region, it suffers from several major limitations due to a combination of coverage biases, low capture efficiency, and errors introduced by PCR amplification. For example an indel located near the end of a target region may not be well covered by sequencing reads, which limits detection ability. Also, the exome capture kits are typically designed to pull down a region of about 400bp around an exon, which can limit detection of large indels within coding regions or near splice sites. On the other hand, albeit with higher cost, WGS comes with several significant benefits, including more uniform coverage, freedom from capture efficiency biases, and the inclusion of the non-coding genome. In the context of detecting indels, it has been shown that the accuracy of indel detection with WGS data is much greater than WES data even within the targeted regions [18] **Table 1** shows that a much higher validation rate of WGS-specific indels, compared to WES-specific indels (84% vs. 57%). Specifically, WGS has a unique advantage over WES in identifying many more indels longer than 5 bp (25 vs. 1). When using WGS, it was estimated that 60X depth of coverage from the HiSeq platform would be needed to recover 95% of the indels detected by Scalpel. In particular, detecting heterozygous indels naturally requires deeper sequencing coverage relative to homozygous indels. **Figure 4**). WGS at 30X using the HiSeq platform is not sufficient for sensitive indel discovery, resulting in at least 25% false negative rates for heterozygous indels. But these requirements can rapidly change with the longer reads and lower error rates provided by newer instruments.

**Figure 4.**
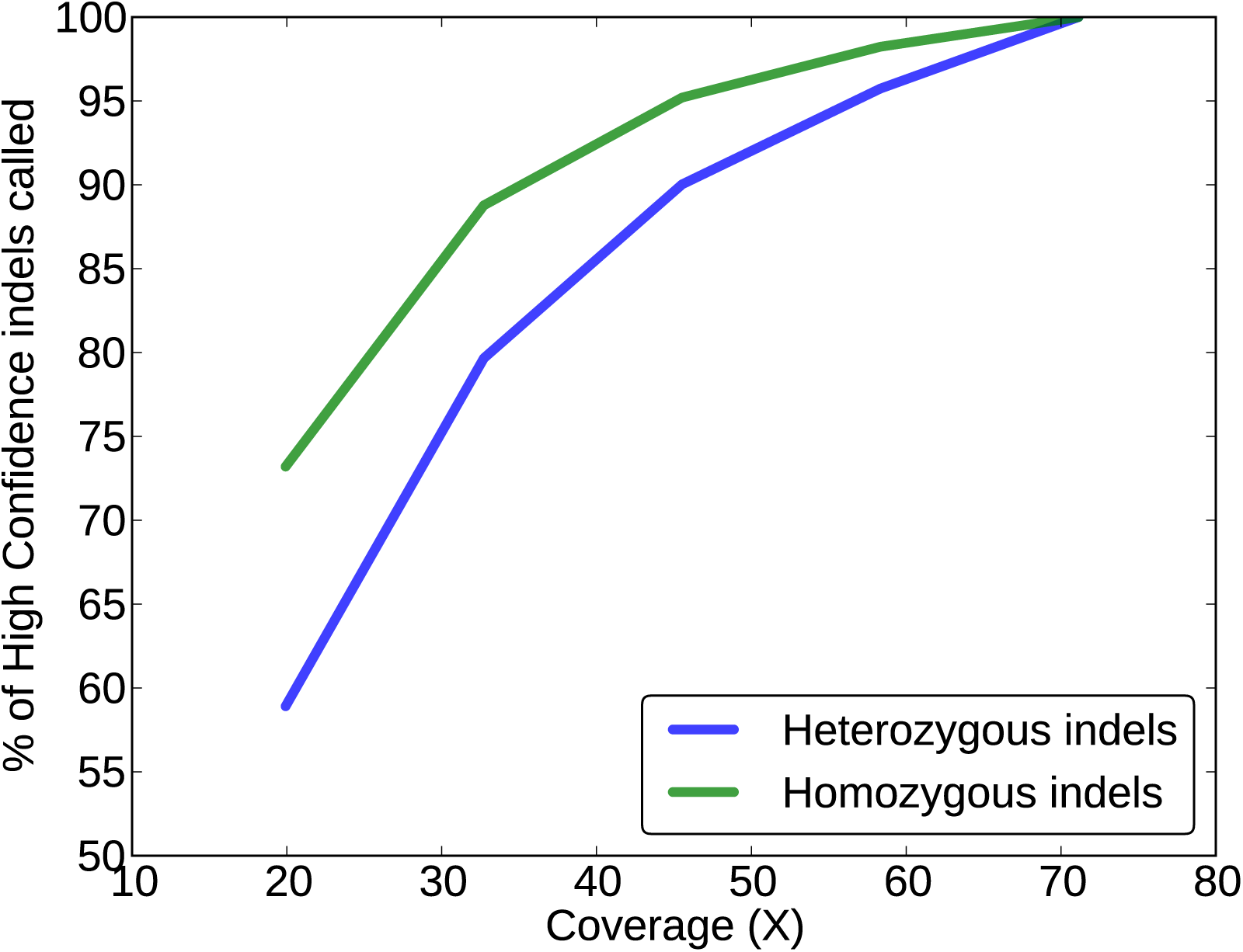
Sensitivity performance of indel detection with WGS data using Scalpel at different coverage on Illumina HiSeq2000 platform. The sensitivity performance is assessed using the high confidence call set shared by WGS and WES data from the same samples (n=8). The Y-axis represents the mean percentage of the high confidence indels revealed at a lower coverage. The X-axis represents the mean coverage of the eight down-sampled genomes. Among the entire call set, about 61% of the indels are heterozygous and the remaining 39% are homozygous. Performance of heterozygous (blue) and homozygous (green) indel detection are shown separately.

### PCR-free protocols

PCR is a widely used and useful technique to amplify DNA fragments of interest and for attaching various linkers or barcodes for sequencing. However, small amounts of contaminating material can also be amplified without discrimination. Also, PCR amplification introduces errors during the library construction step, especially in regions near STRs such as homopolymer A or T runs. These types of errors are due to replication slippage events and result in high variability in the number of repeat elements (**Figure 5**). It becomes then very difficult to distinguish true events at these loci from stutter errors. Moreover, as described in **Box 1**, candidate mutations within STRs can have an ambiguous signature. So for indel analysis, we recommend using PCR-free protocols, which can significantly reduce the number of errors around those loci. Moreover, as reported in this protocol, filtering based on the combination of alternative allele coverage and *k*-mer x^2^ score is an effective strategy to filter out additional false-positives without sacrificing much sensitivity.

**Figure 5.**
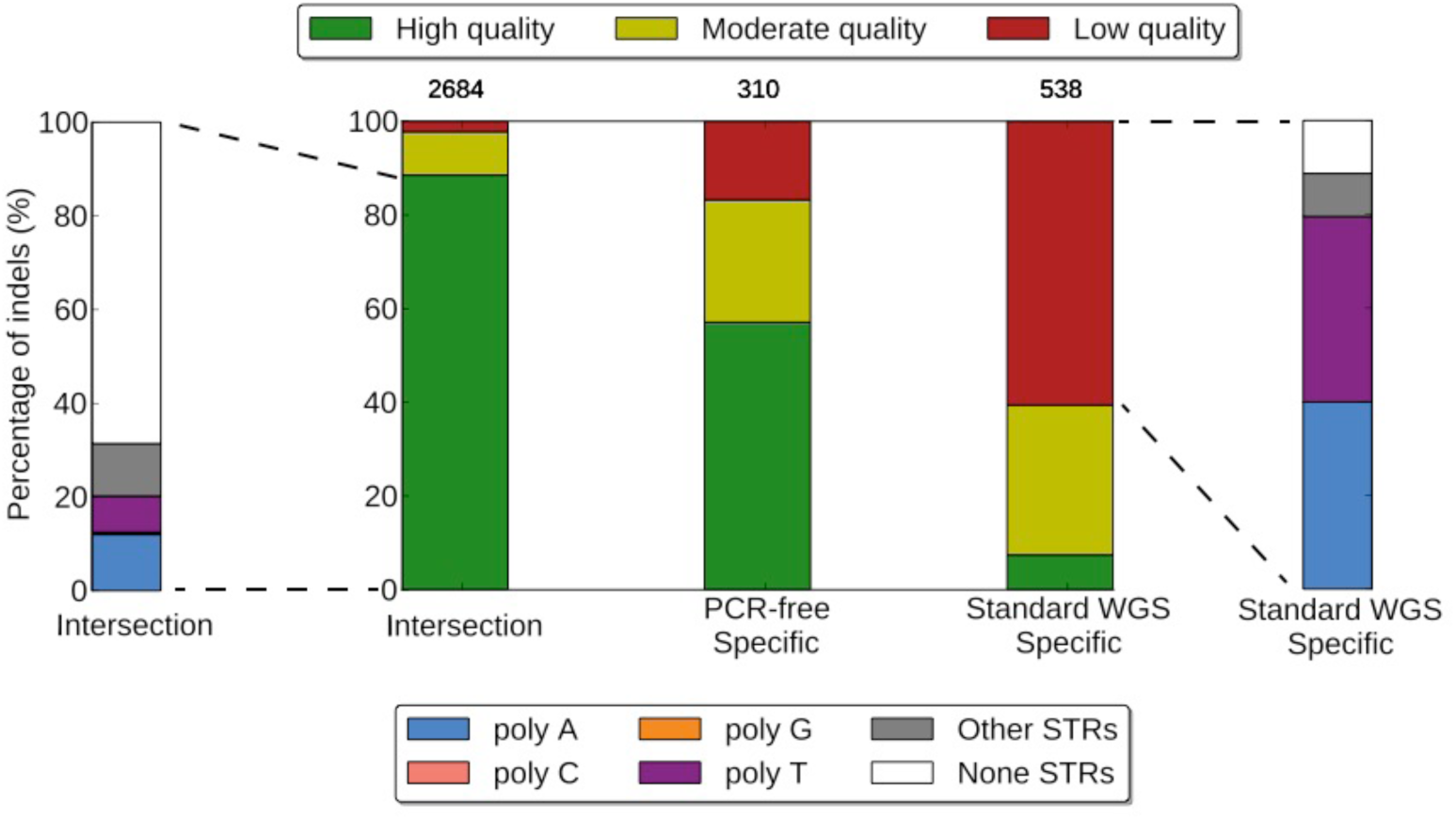
Comparison of standard WGS and PCR-free data based on indel quality. “Intersection” represents the shared indels from both the PCR-free and standard WGS INDELs. The number reported above a call set represent the total number of indels in that call set. Indel calls are further categorized (side-bars) based on their sequence composition: poly-A, poly-C, poly-G, poly-T, other-STR, and non-STR.

### Population studies

Large-scale sequencing studies, involving hundreds or thousands of samples, are now becoming more and more widespread. Here we aim to introduce some of the advantages of having access to a collection of sequenced individuals. Even though Scalpel does not directly provide an API for joint calling on more than four samples, we provide examples and a recommendation on how to take advantage of such information if available. The basic idea is to aggregate all the genetic variants detected in the samples into a database framework with associated genotypes and genomic annotation. There are existing flexible systems for exploring genetic variation for disease and population genetics, such as GEMINI [35]. Analyzing the genetic code of a large cohort of individuals has the potential to shed light on the underpinning mechanisms of complex diseases such as autism and schizophrenia. These studies are generally focused on the detection and analysis of rare variant that can explain the phenotype of the affected individuals.

The population frequency of such rare mutations is usually so low that it is obscured by the noise in the sequencing data, making any real biological signal undetectable. In these circumstances the population can be used to devise effective filtering strategies. For example, in a large-scale autism study where Scalpel was employed [6], the population database was used to identify rare variants by filtering highly polymorphic loci with many more mutations than expected in the general population as well as common variants using minor allele frequency (MAF) cutoffs. Typically, variants for which the minor allele is present in a population above 1% are considered common. By removing these locations from the analysis, the biological signal started to emerge: an enrichment of frame-shift *de novo* mutations in the affected child compared to the unaffected sibling. The highly polymorphic regions were later found to enrich for homopolymers and other STRs, which are known to be more susceptible to sequencing errors. In the case of *de novo* studies, it is extremely unlikely that the same mutation is present as de novo in multiple individuals; in this case the population information can be used again to filter out these candidates as artifacts in the sequencing.

### Cancer studies

Detection of somatic variation in tumor-normal matched samples is complicated by different factors such as ploidy, clonality, and purity of the input material. Moreover, the sensitivity and specificity of any somatic mutation calling approach varies along the genome due to differences in sequencing read depths, error rates, variant allele fractions (VAF) of mutations, etc. Accounting for all these variables poses a very complex and challenging problem. However, the proper filtering parameters can eliminate the majority of Scalpel’s false-positive calls. For example, **Figure 6 and 7** show the effects of different phred-scaled Fisher’s exact score cutoffs used for filtering on a pair of highly concordant primary and metastatic samples from Branon et.al [36]. Figure 7 demonstrates that indels with a phred-scaled Fisher’s exact score below 10 tend to have low VAF and are much more likely to be sequencing errors. In fact, the allele fraction of mutations exclusive to either the primary tumor or the metastasis is significantly lower with higher (more stringent) cutoffs. Similarly, the VAF distribution of the indels found only in the primary tumor shifts towards the expected distribution (with a peak at ~20%) as more conservative Fisher’s exact test cutoffs are used. Not all errors are eliminated though, especially in regions where very low support for a mutation in the normal or the tumor precludes the assembly of the reads. We are actively researching enhanced algorithms for such regions, including using a joint assembly within the same de Bruijn graph of the reads from both the tumor and the normal samples.

**Figure 6.**
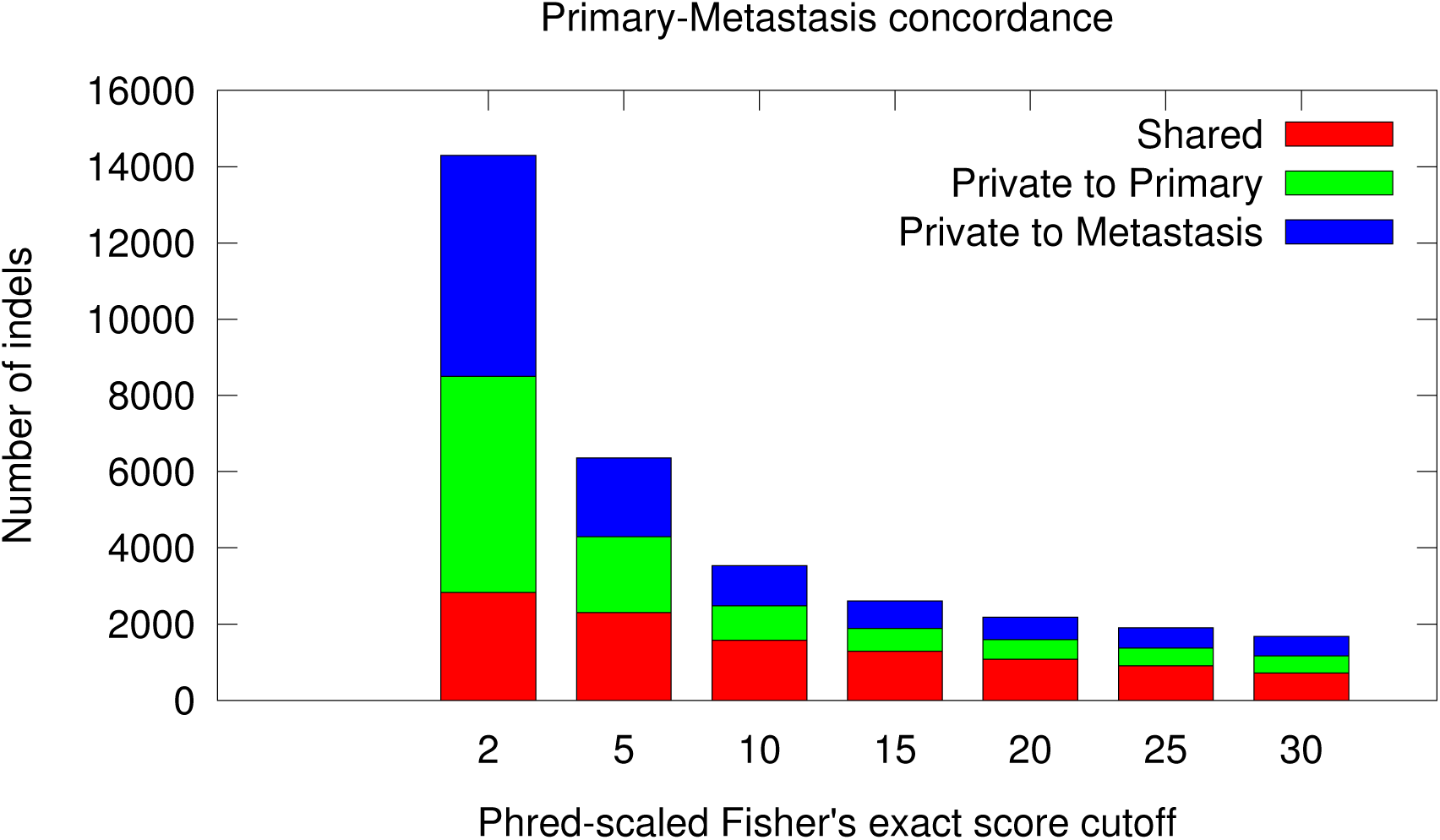
Whole genome mutational concordance. Concordance and discordant indel mutations as a function of the phred-scaled Fisher’s exact score cutoff between primary and metastasis for a pair of highly concordant colorectal cancer samples.

**Figure 7.**
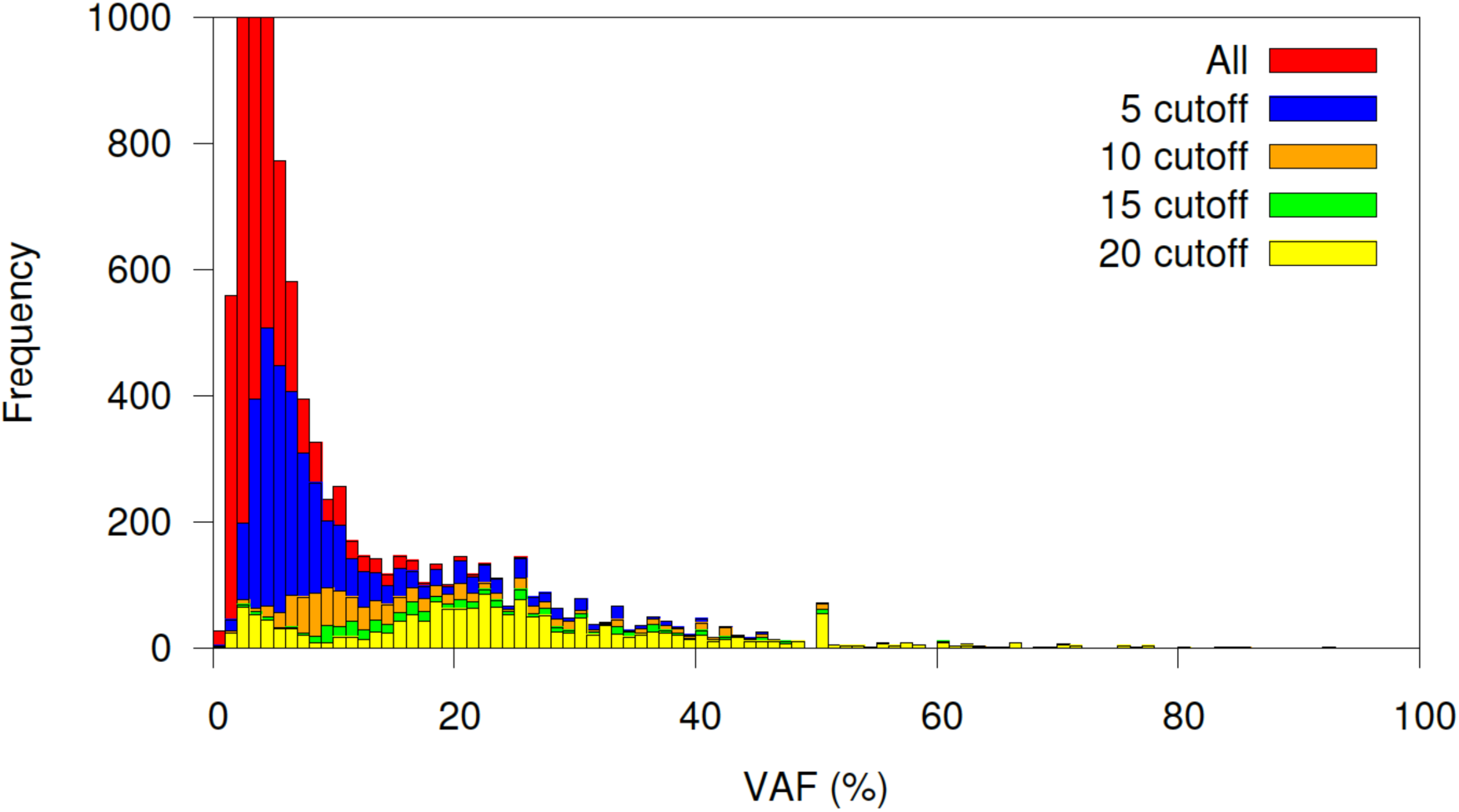
Variant allele fraction (VAF) distribution. Distribution of VAF as a function of different phred-scaled Fisher’s exact score cutoffs for the somatic indels detected in the primary tumor (from **Figure 6**).

## MATERIALS

### EQUIPMENT

Δ CRITICAL Make sure that the listed software tools are available within your UNIX PATH setting. For example, if you have your tools installed in a ‘/path/to/your/tools’ directory, you can update your PATH setting to include this path by using the following command:

~~~
% export PATH=/path/to/your/tools:$PATH
You can also add the command to your UNIX setting file ~/.bashrc.
~~~

– BWA [34] version 0.7.12 (http://bio-bwa.sourceforge.net/)
– Samtools [21] version 1.2 (http://samtools.sourceforge.net/)
– bcftools version 1.2 (http://samtools.github.io/bcftools/)
– picard version 1.130 (broadinstitute.github.io/picard/)
– Scalpel version 0.4.1 (scalpel.sourceforge.net)
– Bedtools [39] version 2.23.0 (https://github.com/arq5x/bedtools2)
– PyVCF version 0.6.7 (https://github.com/jamescasbon/PyVCF)
– ANNOVAR [38] version 2015-03-22 (http://annovar.openbioinformatics.org/)
– R version 2.15 (http://www.r-project.org)
– Gnuplot version 4.4 (http://www.gnuplot.info/)
– IGV [40] version 2.3 (https://www.broadinstitute.org/igv/)
– Download the scripts provided for visualizing and quality control of the indels available in Scalpel’s resource bundle.

### EQUIPMENT SETUP

#### Hardware setup

The software used in this protocol is intended for operation on a 64-bit machine, running a 64-bit version of the Linux operating system. We recommend using a machine with at least 1.2Tb of disk storage available for whole genome analysis and a minimum of 64 GB of RAM. The software will scale to the number of cores available, and recommend at least ten cores if possible, especially for whole genome analysis.

#### Software setup

Download relevant files host on the Scalpel website, including the resource bundle:

~~~
% wget --no-check
http://sourceforge.net/projects/scalpel/files/scalpel–0.5.1.tar.gz; tar zxvf scalpel-0.5.1.tar.gz; cd scalpel–0.5.1; make; export PATH=../scalpel–0.5.1:$PATH
% tar zxvf protocol_bundle-0.5.1.tar.gz; cd protocol_bundle–0.5.1
~~~

Download tools that are required for the indel analysis, including bwa, samtools, bcftools, picard, scalpel, vt, bedtools, ANNOVAR (needs registration):

~~~
% wget --no-check http://sourceforge.net/projects/bio-bwa/files/bwa-0.7.12.tar.bz2; tar jxf bwa-0.7.12.tar.bz2; cd bwa- 0.7.12; make; cd ../; export PATH=./bwa-0.7.12:$PATH
% wget --no-check
http://sourceforge.net/projects/samtools/files/samtools/1.2/samtools-1.2.tar.bz2; tar jxf samtools-1.2.tar.bz2; cd samtools-1.2; make; cd ..; export PATH=./samtools-1.2:$PATH
% wget -no-check
https://github.com/samtools/bcftools/releases/download/1.2/bcftools-1.2.tar.bz2; tar bcftools-1.2.tar.bz2; cd bcftools-1.2; make; cd ..; export PATH=./bcftools-1.2:$PATH
% wget --no-check
https://github.com/broadinstitute/picard/releases/download/1.130/picard-tools-1.130.zip; unzip picard-tools-1.130.zip; export PATH=./picard-tools-1.130:$PATH
% wget --no-check
https://github.com/arq5x/bedtools2/releases/download/v2.23.0/bedtools-2.23.0.tar.gz; tar zxvf bedtools-2.2 3.0.tar.gz;cd bedtools2; make; cd ..; export PATH=./bedtools-2.2 3.0/bin:$PATH
% wget
http://www.openbioinformatics.org/annovar/download/register-for-download/annovar.latest.tar.gz --no-check-certificate; tar zxvf annovar.latest.tar.gz; export PATH=./annovar:$PATH
% git clone https://github.com/jamescasbon/PyVCF.git cd PyVCF; python setup.py install; cd ..
~~~

### PROCEDURE

**Download the example sequencing data and reference** | Time: ~6 hours This protocol and bioinformatics software is generally applicable and optimized for Illumina NGS data, including WGS and exome captured sequencing data. We use the publicly WGS data on the family of NA12878 as an illustration in this protocol, but some specific parameters like filtering criterion need to be adjusted accordingly.) We also have additional boxes for expanding analysis to somatic (e.g., tumor/normal) and singlesampling indel calling.

**1|** Download the example sequencing reads of the Hapmap quad family from the Illumina Platinum Genome project (*_1*fastq.gz and *_2*fastq.gz denote paired end reads):

~~~
% wget --no-check
ftp://ftp.sra.ebi.ac.uk/vol1/fastq/ERR194/ERR194146/ERR194146_1.fastq.gz
% wget --no-check
ftp://ftp.sra.ebi.ac.uk/vol1/fastq/ERR194/ERR194146/ERR194146_2.fastq.gz
% wget --no-check
ftp://ftp.sra.ebi.ac.uk/vol1/fastq/ERR194/ERR194147/ERR194147_1.fastq.gz
% wget --no-check
ftp://ftp.sra.ebi.ac.uk/vol1/fastq/ERR194/ERR194147/ERR194147_2.fastq.gz
% wget --no-check
ftp://ftp.sra.ebi.ac.uk/vol1/fastq/ERR194/ERR194151/ERR194151_1.fastq.gz
% wget --no-check
ftp://ftp.sra.ebi.ac.uk/vol1/fastq/ERR194/ERR194151/ERR194151_2.fastq.gz
% wget --no-check
ftp://ftp.sra.ebi.ac.uk/vol1/fastq/ERR324/ERR324432/ERR324432_1.fastq.gz
% wget --no-check
ftp://ftp.sra.ebi.ac.uk/vol1/fastq/ERR324/ERR324432/ERR324432_2.fastq.gz
~~~

**2|** Download the human reference genome hg19:

~~~
% wget --no-check
http://hgdownload.cse.ucsc.edu/goldenPath/hg19/bigZips/hg19.2bit
% wget --no-check
http://hgdownload.cse.ucsc.edu/admin/exe/linux.x86_64/twoBitToFa
~~~

**3|** Convert the *.2bit genome to *.fa format and index it with bwa (Note you can also download the fasta file directly, although may take much longer):

~~~
% chmod +x twoBitToFa; ./twoBitToFa hg19.2bit hg19.fa
% bwa index hg19.fa
~~~

**Align the NGS reads to the genome** | Time: ~40 hours

**4|** Align reads to reference for each sample separately with bwa mem:

~~~
% bwa mem -t 10 -R ‘@RG\tID:NA12877\tSM:NA12877’ hg19.fa ERR194146_1.fastq.gz ERR19414 6_2.fastq.gz | samtools view -h -S - b > NA12 8 77.bam
% bwa mem -t 10 -R ‘@RG\tID:NA12878\tSM:NA12878’ hg19.fa ERR194147_1.fastq.gz ERR19414 7_2.fastq.gz | samtools view -h -S - b > NA12 8 78.bam
% bwa mem -t 10 -R ‘@RG\tID:NA12881\tSM:NA12881’ hg19.fa
ERR32 44 32_1.fastq.gz ERR324432_2.fastq.gz | samtools view -h -S - b > NA12 8 81.bam
% bwa mem -t 10 -R ‘@RG\tID:NA12882\tSM:NA12882’ hg19.fa ERR194151_1.fastq.gz ERR194151_2.fastq.gz | samtools view -h -S - b > NA12 8 82.bam
~~~

**5|** Sort the bam files by chromosome coordinates with samtools:

~~~
% samtools sort -m 4G NA12877.bam NA12877.sort
% samtools sort -m 4G NA12878.bam NA12878.sort
% samtools sort -m 4G NA12881.bam NA12881.sort
% samtools sort -m 4G NA12882.bam NA12882.sort
% rm -f NA12 8 77.bam NA12 8 78.bam NA12 8 81.bam NA12 8 82.bam
~~~

**6|** Mark duplicated reads within the alignment with picard tools:

~~~
% java -jar -Xmx10g picard MarkDuplicates INPUT=NA12 8 77.sort.bam OUTPUT=NA12877.sort.markdup.bam METRICS_FILE=NA12877.sort.metric
% java -jar -Xmx10g picard MarkDuplicates INPUT=NA12 8 78.sort.bam
OUTPUT=NA12878.sort.markdup.bam METRICS_FILE=NA12878.sort.metric
% java -jar -Xmx10g picard MarkDuplicates INPUT=NA12881.sort.bam
OUTPUT=NA12 881.sort.markdup.bam METRICS_FILE=NA12 881.sort.metric
% java -jar -Xmx10g picard MarkDuplicates INPUT=NA12882.sort.bam
OUTPUT=NA12 882.sort.markdup.bam METRICS_FILE=NA12 882.sort.metric
% rm -f NA12877.sort.bam NA12878.sort.bam NA12881.sort.bam NA12 8 82.sort.bam
~~~

**7|** Perform a basic quality control of the alignment files with samtools:

~~~
% samtools flagstat NA12877.sort.markdup.bam >
NA12877.sort.markdup.bam.simplestats
% samtools flagstat NA12878.sort.markdup.bam >
NA12878.sort.markdup.bam.simplestats
% samtools flagstat NA12881.sort.markdup.bam >
NA12881.sort.markdup.bam.simplestats
% samtools flagstat NA12882.sort.markdup.bam >
NA12882.sort.markdup.bam.simplestats
~~~

Δ CRITICAL: In order to generate reliable indel calls, accurate alignment of the NGS short reads are of great importance. If the DNA is derived from blood sample, the mapping rate of Illumina Hiseq reads is typically higher than 90%. Lower mapping rates indicate either contaminations of DNA from other species (e.g. bacterial DNA from saliva samples) or poor quality of the sequencing experiments. In addition, excessive numbers of duplicated reads are usually due to issues with library construction and PCR amplification. Table 2 lists the number of reads generated for each sample and the reads mapped to the human genome hg19.

**Table 2a.**
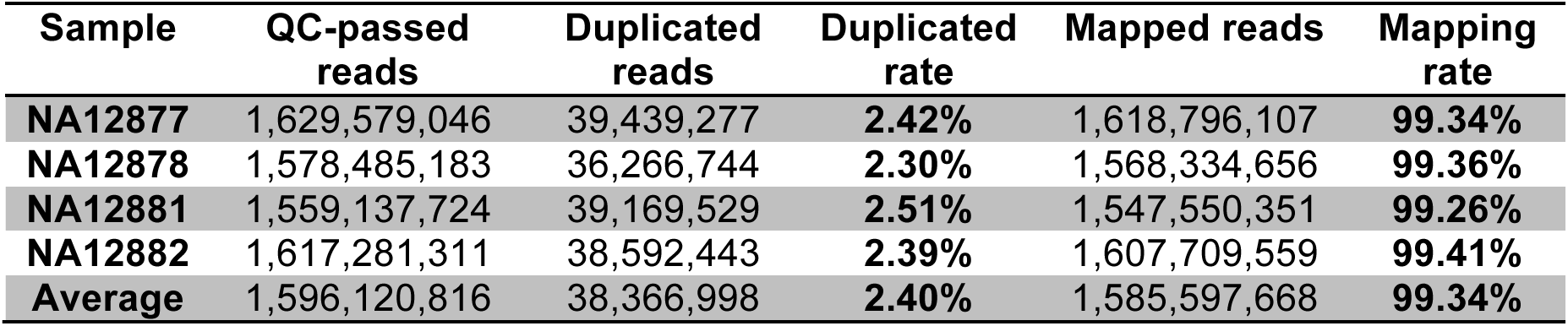
Expected QC-passed read and mapping statistics

| Sample | QC-passed reads | Duplicated reads | Duplicated rate | Mapped reads | Mapping rate |
| --- | --- | --- | --- | --- | --- |
| NA12877 | 1,629,579,046 | 39,439,277 | 2.42% | 1,618,796,107 | 99.34% |
| NA12878 | 1,578,485,183 | 36,266,744 | 2.30% | 1,568,334,656 | 99.36% |
| NA12881 | 1,559,137,724 | 39,169,529 | 2.51% | 1,547,550,351 | 99.26% |
| NA12882 | 1,617,281,311 | 38,592,443 | 2.39% | 1,607,709,559 | 99.41% |
| Average | 1,596,120,816 | 38,366,998 | 2.40% | 1,585,597,668 | 99.34% |

**Perform indel variant calling and downstream filtering** | Time: ~8 hours

8| Run Scalpel in the “de novo” mode to perform multi-sample calling for a family. In this example, we use NA12882 as the affected individual. The NA12881 is the unaffected individual accordingly:

~~~
% scalpel-discovery --denovo --dad NA12877.sort.markdup.bam –mom
NA12 8 78.sort.markdup.bam –aff NA12 882.sort.markdup.bam –sib
NA12881.sort.markdup.bam --bed
SeqCap_EZ_Exome_v3_primary.scalpel.bed --ref hg19.fa --numprocs 10 --two-pass
~~~

Δ CRITICAL: In both “denovo” and “somatic” mode, Scalpel is optimized to achieve high sensitivity, but may include some false positives. To control for this, we recommend using the --two-pass option in Scalpel, which undergoes a second round of indel verification to reduce the likely false calls.

**9|** Export the inherited and denovo mutations from the Scalpel database (in target only):

~~~
% scalpel-export --denovo --db outdir/main/inherited.db –bed
SeqCap_EZ_Exome_v3_primary.scalpel.bed --ref hg19.fa --intarget - -min-alt-count-affected 10 --max-chi2-score 10.8 > inherited.onepass.vcf
% scalpel-export --denovo --db outdir/twopass/denovos.db –bed
SeqCap_EZ_Exome_v3_primary.scalpel.bed --ref hg19.fa --intarget - -min-alt-count-affected 10 --max-chi2-score 10.8 --min-coverage- unaffected 20 > denovo.twopass.vcf
~~~

**10|** Identify and mark indels within STR regions using ms-detector:

~~~
% sh ./msdetector/msdetector.sh -r 50 -d 2 -g hg19.fa –i
inherited.onepass.vcf > inherited.onepass.vcf.ms
% sh ./msdetector/msdetector.sh -r 50 -d 2 -g hg19.fa –i
denovo.twopass.vcf > denovo.twopass.vcf.ms
~~~

**11|** Save indels within and outside STR regions into different vcf files:

~~~
% awk -F “\t” ‘{if($0 ~ /^#/){print $0} else{if($16==“yes”)
print} }’ inherited.onepass.vcf.ms | cut -f1-13 >
inherited.onepass.vcf.ms.in
% awk -F “\t” ‘{if($0 ~ /^#/){print $0} else{if($16==“no”) print}
}’ inherited.onepass.vcf.ms | cut -f1-13 >
inherited.onepass.vcf.ms.out
% awk -F “\t” ‘{if($0 ~ /^#/){print $0} else{if($16==“yes”)
print} }’ denovo.twopass.vcf.ms | cut -f1-13 >
denovo.twopass.vcf.ms.in
% awk -F “\t” ‘{if($0 ~ /^#/){print $0} else{if($16==“no”) print}
}’ denovo.twopass.vcf.ms | cut -f1-13 >
denovo.twopass.vcf.ms.out
~~~

Δ CRITICAL: Low quality indel calls (potential false-positives) are usually found within low coverage regions, or have an unbalanced number of reads supported the alternative allele.

**12|** Filter out false positive calls by adjusting coverage and/or chi-squared thresholds for your data:

~~~
% awk -F “\t” ‘{if($0 ~ /^#/){print $0} else {if(!
($7~/LowAltCntAff/ && $7~/HighChi2score/) ) print} }’
inherited.onepass.vcf.ms.out > inherited.onepass.vcf.ms.out.hq
% awk -F “\t” ‘{if($0 ~ /^#/){print $0} else {if(!
($7~/LowAltCntAff/ || $7~/HighChi2score/ || $7~/LowCovUnaff/) )
print} }’ denovo.twopass.vcf.ms.out >
denovo.twopass.vcf.ms.out.hq
~~~

**13|** (Optional) Perform additional filtering of the de novo calls using the python script provided in the Scalpel resource bundle. This script supports filtering indels by alternative allele coverage (aac), chi-square scores (chi), and parental coverage (pc):

~~~
% python denovo-multi-filter.py -i denovo.twopass.vcf.ms.out -f NA12877 -m NA12878 -a NA12882 -u NA12881 -aac 10 -chi 10.8 -pc 20 -o denovo.twopass.vcf.ms.out.filter
~~~

**14|** (Optional) Extract a subset of indels based on other annotation using bedtools:

~~~
% bedtools intersect -wa -u -a inherited.onepass.vcf.ms.out.hq –b
clinvar_main.bed > inherited.onepass.vcf.ms.out.hq.clinvar
~~~

**15|** Summarize indel calls with histogram of mutations by size:

~~~
% grep -v “#” inherited.onepass.vcf.ms.out.hq
denovo.twopass.vcf.ms.out.hq | awk ‘{print length($5)-
length($4)}’ > all.indel.size.txt % gnuplot44 -e “outfile=‘indel_size_dist.pdf’;
infile=‘all.indel.size.txt’” size_dist.gnu
~~~

**Figure 9.**
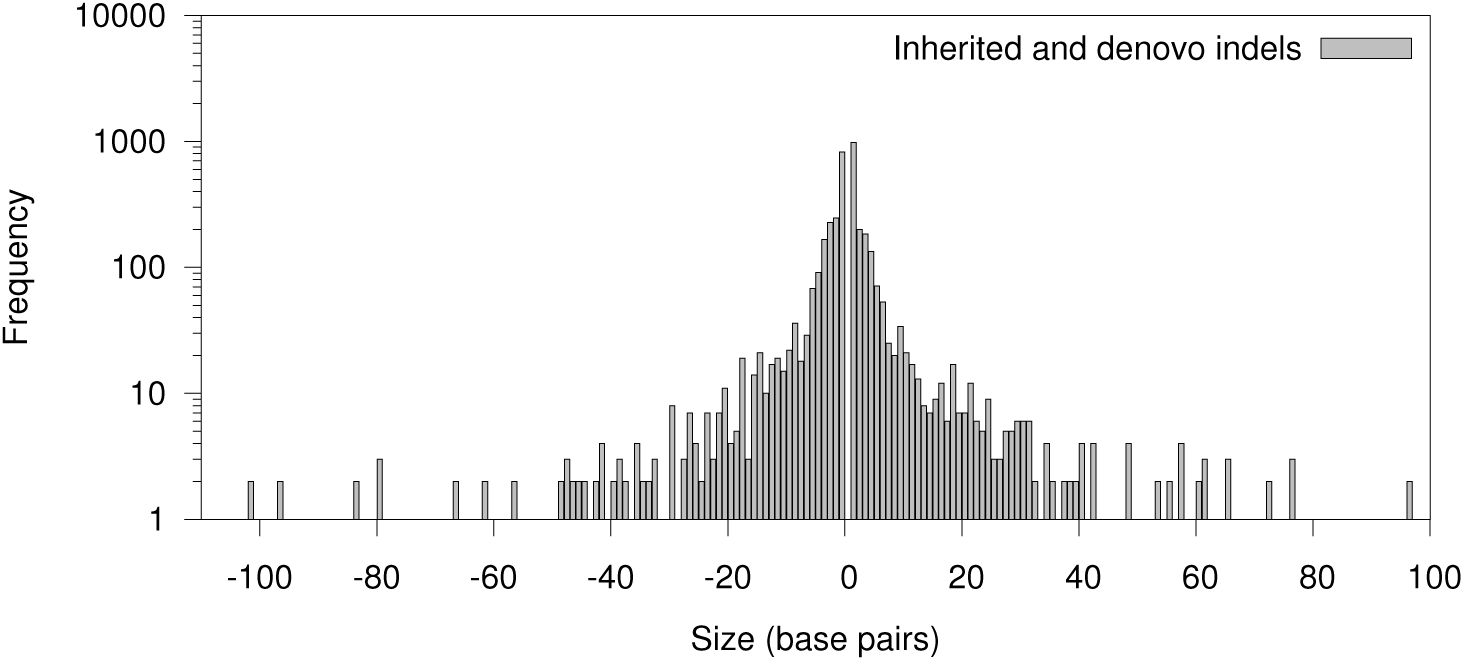
Size distribution of inherited and denovo indels. We should expect a lognormal distribution of indels with majority of them being short, i.e. less than 5bp.

16| Characterize low quality homopolymer indels calls with histogram of mutations by VAF:

~~~
% cat denovo.twopass.vcf.ms inherited.onepass.vcf.ms | grep –v
‘#’ |grep ‘yes’ | awk -F “\t” ‘{if( ($7~/LowAltCntAff/ &&
$7~/HighChi2score/) || $7~/LowCovUnaff/ ) print}’ > combine.\ ms.txt
% for i in A C G T; do awk -v j=$i ‘$0!~/^#/ { if($15==j) {
split($12,a,”:”); if(a[1]==“0/1” || a[1]==“1/1”)
split(a[2],b,”,”); print b[1] “\t” b[2]} }’ combine.ms.txt >
poly${i}.VAF.txt; done
% gnuplot44 -e “outfile=‘homo.vaf.pdf’; infileA=‘polyA.VAF.txt’;
infileC=‘polyC.VAF.txt’; infileG=‘polyG.VAF.txt’;
infileT=‘polyT.VAF.txt’ “ hp.vafdist.gnu
~~~

Δ CRITICAL: There are usually much higher sequencing biases in GC-extreme regions. Indels within STRs especially homopolymer A or T runs are major source of false positive variant calls.

**Figure 10.**
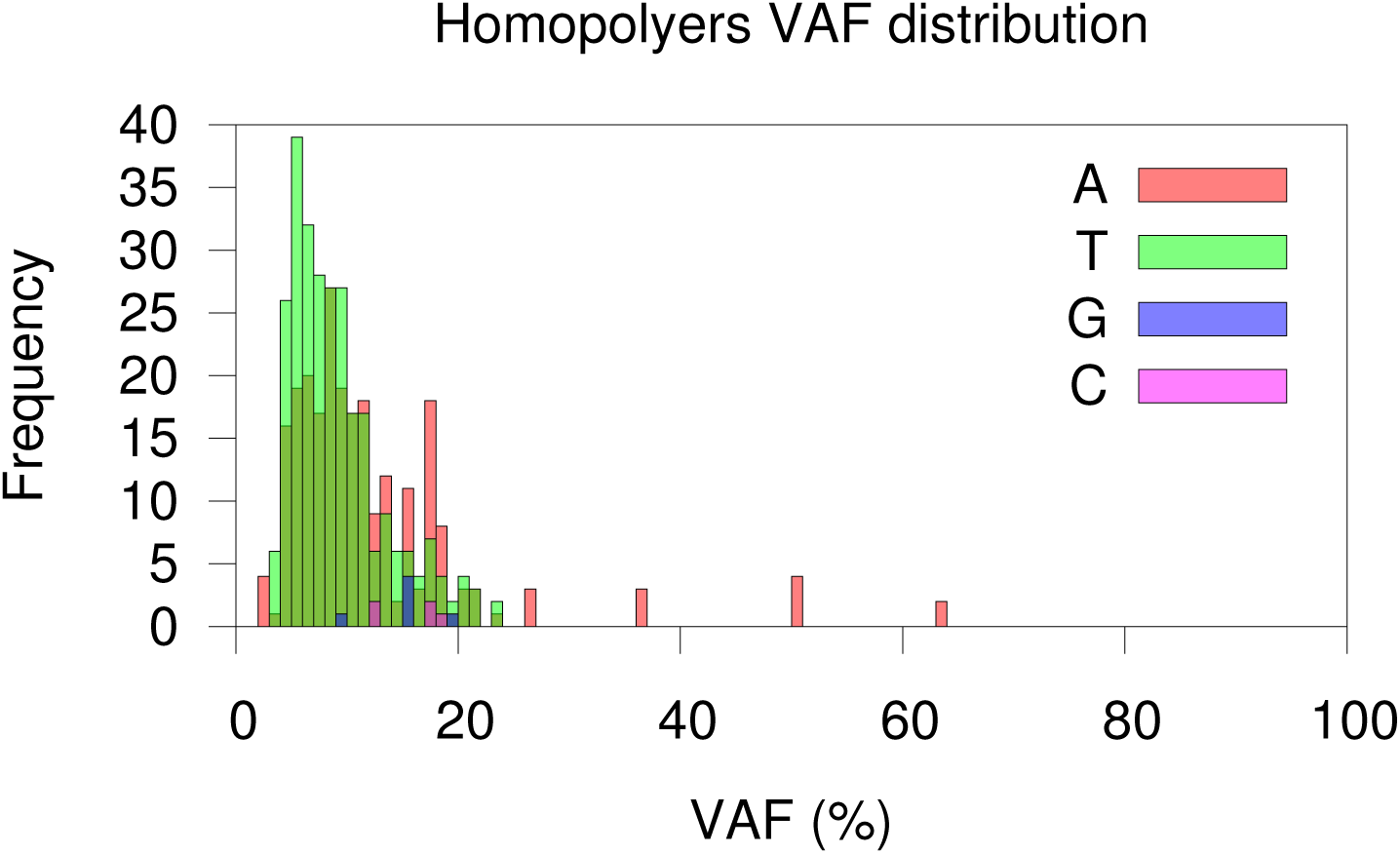
Histograms of low quality homopolymer indels by category. Homopolymer A or T indels should be more abundant than C or G indels in the call set, especially indels with low variant allele fraction (VAF). Due to the limitation of PCR amplification, homopolymer A or T runs are more like result in inaccurate molecules [18].

**17|** Summarize inherited indels with variant allele fractions (VAF %):

~~~
% awk -F’\t’ ‘$0!~/^#/ {split($12,a,”:”); if(a[1]==“0/1” ||
a[1]==“1/1”) split(a[2],b,”,”); print b[1] “\t” b[2]}’
inherited.onepass.vcf.ms.out > inherited.onepass.vcf.ms.out.vaf
% awk -F’\t’ ‘$0!~/~#/ {split($12,a,”:”); if(a[1]==“0/1” ||
a[1]==“1/1”) split(a[2],b,”,”); print b[1] “\t” b[2]}’
inherited.onepass.vcf.ms.out.hq >
inherited.onepass.vcf.ms.out.hq.vaf
% gnuplot44 -e “outfile=‘inherited.VAFdist.pdf’;
infileAll=‘inherited.onepass.vcf.ms.out.vaf’;
infileHq=‘inherited.onepass.vcf.ms.out.hq.vaf’”
vafdistplot.inherited.qual.gnu
~~~

Δ CRITICAL: The filtering cascade should not reduce the sensitivity of inherited indels by a lot. One should expect a relatively balanced number of reads support each inherited indel, indicating high confidence for these calls.

**Figure 11.**
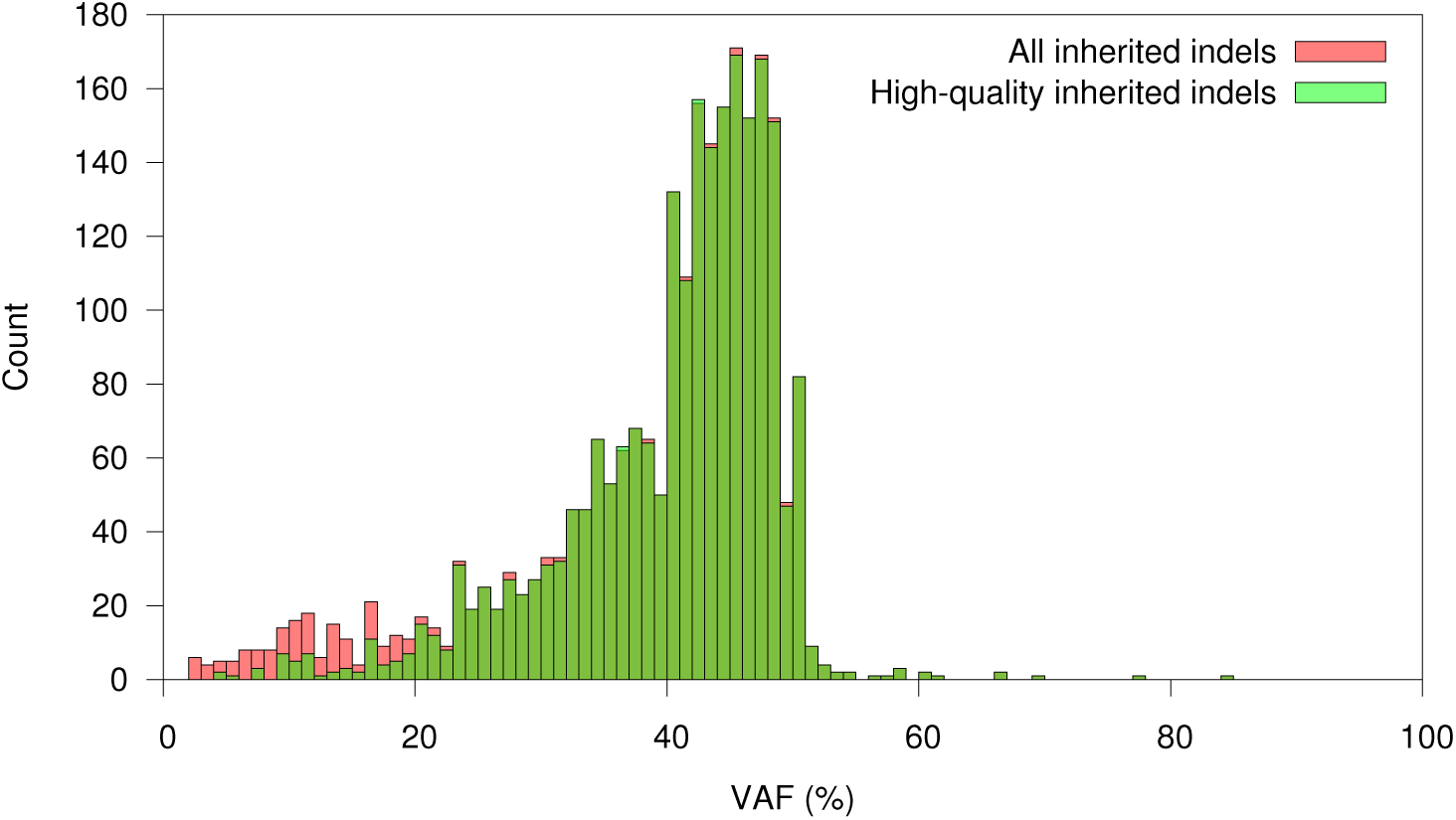
Variant allele fractions (VAF %) of the inherited indels. VAF of inherited indels should approximately follow a normal distribution with a mean of about 50%. In practice, due to sequencing and alignment biases, the mean of the normal distribution is usually slightly smaller than 50%.

**18|** Determine the number of indels remained after each step of the filtering:

~~~
% for i in *.vcf.* ; do echo $i; grep -v “#” $i | wc -l;done >
indel.count.txt
~~~

**19|** Split the multi-sample VCF to an individual file for NA12882:

~~~
% for file in *.hq; do bgzip -c $file > $file.gz ; tabix -p vcf
$file.gz; done
% for file in *.hq.gz; do bcftools view -c1 -Oz -s NA12882 –o
${file/.gz*/.$sample.vcf.gz} ${file}; gunzip
${file/.gz*/.$sample.vcf.gz}; done
~~~

**20|** Filter the single VCF files based on Chi-Square score and allele coverage

~~~
% python single-vcf-filter.py -i
inherited.onepass.vcf.ms.out.hq.NA12 8 82.vcf -mc 10 -chi 10.8 o
inherited.onepass.vcf.ms.out.hq.NA12 8 82.filter.vcf
% python single-vcf-filter.py -i
denovo.twopass.vcf.ms.out.hq.NA12882.vcf -mc 10 -chi 10.8 o
denovo.twopass.vcf.ms.out.hq.NA12 882.filter.vcf
~~~

Annotation and visualization of the indel calls | Time: <5 min

21| Prepare and create the input format required by ANNOVAR:

~~~
% annovar=/path-to-annovar/
% $annovar/convert2annovar.pl -format vcf4
inherited.onepass.vcf.ms.out.hq.NA12 8 82.filter.vcf >
inherited.onepass.vcf.ms.out.hq.NA12 8 82.filter.vcf.avinput
% $annovar/convert2annovar.p1 -format vcf4
denovo.twopass.vcf.ms.out.hq.NA12 882.filter.vcf >
denovo.twopass.vcf.ms.out.hq.NA12 882.filter.vcf.avinput
~~~

**22|** Annotate and intersect indels with gene regions using ANNOVAR:

~~~
% $annovar/annotate_variation.pl -buildver hg19
inherited.onepass.vcf.ms.out.hq.NA12 8 82.filter.vcf.avinput
$annovar/humandb
% $annovar/annotate_variation.pl -buildver hg19
denovo.twopass.vcf.ms.out.hq.NA12 882.filter.vcf.avinput
$annovar/humandb
~~~

**23|** Summarize coding region indels by size in R:

~~~
% cat
inherited.onepass.vcf.ms.out.hq.NA12 8 82.vcf.avinput.exonic_varian
t_function | egrep -v ‘unknown|stopgain’ | cut -f 2,7,8 | cut –d
“ “ -f 2 | awk ‘{if($2==“-”) print $1”\t”length($3);else if
($3==“-”) print $1”\t”length($2)}’ > type_and_size.txt
% R
> indel=read.table(“type_and_size.txt”, header=FALSE)
> colnames(indel)= c(“type”,”size”)
> indel_30=indel[indel[,2]<=30,]
> indel.table <-
table(indel_30$type,factor(indel_30$size,lev=1:30) )
> pdf(‘indelsize_by_type.pdf’, width=12, height=8)
> barplot(indel.table, main=“indel distribution within coding
sequence (CDS)”, xlab=““, col=c(“green”,”red”), legend =
rownames(indel.table))
> dev.off()
~~~

**24|** Filter the indels based on population allele frequencies:

~~~
% $annovar/annotate_variation.pl -filter –out
inherited.onepass.vcf.ms.out.hq.NA12 8 82.filter –dbtype
popfreq_max_20150413 -build hg19
inherited.onepass.vcf.ms.out.hq.NA12 8 82.filter.vcf.avinput
$annovar/humandb/
% $annovar/annotate_variation.pl -filter –out
denovo.twopass.vcf.ms.out.hq.NA12882.filter –dbtype
popfreq_max_20150413 -build hg19
denovo.twopass.vcf.ms.out.hq.NA12 882.filter.vcf.avinput
$annovar/humandb/
~~~

**25|** Annotate novel indels that were not reported by population database before (1000G, ESP6500, ExAC, CG46):

~~~
% $annovar/annotate_variation.pl -buildver hg19
inherited.onepass.vcf.ms.out.hq.NA12 8 82.filter.hg19_popfreq_max_2
0150413_filtered $annovar/humandb
% $annovar/annotate_variation.pl -buildver hg19
denovo.twopass.vcf.ms.out.hq.NA12 882.filter.hg19_popfreq_max_2015
0413_filtered $annovar/humandb
~~~

**26|** Retrieve frame-shift mutations, which are potentially loss-of-function:

~~~
% awk ‘{if($2==“frameshift”) print}’
inherited.onepass.vcf.ms.out.hq.NA12 8 82.filter.hg19_popfreq_max_2
0150413 filtered.exonic variant function >
inherited.onepass.vcf.ms.out.hq.NA12882.filter.hg19_popfreq_max_2
0150413_filtered.exonic_variant_function.fs.txt
% awk ‘{if($2==“frameshift”) print}’
denovo.twopass.vcf.ms.out.hq.NA12882.filter.hg19_popfreq_max_2015
0413_filtered.exonic_variant_function >
denovo.twopass.vcf.ms.out.hq.NA12882.filter.hg19_popfreq_max_2015
0413_filtered.exonic_variant_function.fs.txt
~~~

### TROUBLESHOOTING

**Table 1a.** Troubleshooting table

| Step | Problem | Possible reason | Solution |
| --- | --- | --- | --- |
| 4-8 | Bash cannot find the command for a tool. | The incorrect directory of the tool is exported into \$PATH. | Make sure the root directory containing the executable files are exported into \$PATH, otherwise just use the absolute path of the tool. |
| 7 | Low mapping rate (<90%) | Untrimmed barcodes on reads, Poor quality reads, contamination, or invalid mapper settings | Evaluate the read quality using FastQC and trim if necessary. Reads that fail to map can be realigned with more sensitive settings. Reads that still fail to map are likely contamination and can be safely ignored. |
| 9 | There are very few inherited indels in the outputs and/or there is an excessive number of de novo indels. | The incorrect database might have been used. | Use the “main” folder for exporting inherited indels and use the “two-pass” folder for exporting de novo indels. |
| 9 | Excessive numbers of de novo indels | Poor quality reads require most stringent filtering | Adjust the minimum coverage and chi squared parameters to reduce the number of false positive calls |
| 11 | The number of variants reduces after running ms-detector. | ms-detector might have introduced blank space into some columns | Make sure you follow the protocol commands and use -F "\t" in the awk script. |
| 14-17 | Gnuplot is not producing the figure. | Some earlier versions of gnuplot do not have the necessary functions. | Use gnuplot version 0.44 or later. |

Following this protocol, it will take ~48 h to finish the analysis of the exonic indels in the example WGS data on a machine with ten processing cores and at least 53 GB of RAM. However, the time could be variable depending on the user’s actual computational power. Most of the running time is spent on read alignment. Notably, we can expect in the near future, there will be studies with larger sample size and deeper sequencing. So the running may take longer, assuming the computer power remains the same.

Step 1-3, download the example data from the public sever: ~6 hours

Steps 4–7, align the WGS reads to the genome: ~40 h

Steps 8–17, perform exonic indel variant calling and downstream filtering: ~5 h

Step 18-20, annotate and visualize of the indel calls: <5 min

### ANTICIPATED RESULTS

Expected number of indels that remains during the filtering cascade:

Since calling de novo indels requires a more sensitive analysis of the family members, we recommend using the --two-pass search option when discovering denovo events. Many more inherited indels will persist through the filtering cascade, relative to the number of de novo events. This is because de novo events are extremely rare in comparison to inherited indels. De novo mutations are also particularly vulnerable to batch effects and random errors, as a correct analysis requires both high sensitivity and specificity in the entire family. In fact, among the in target indels, about 51% of the inherited ones are of high quality while only 5% of the denovo ones survived the filtering cascade (**Figure 12**).

**Figure 12.**
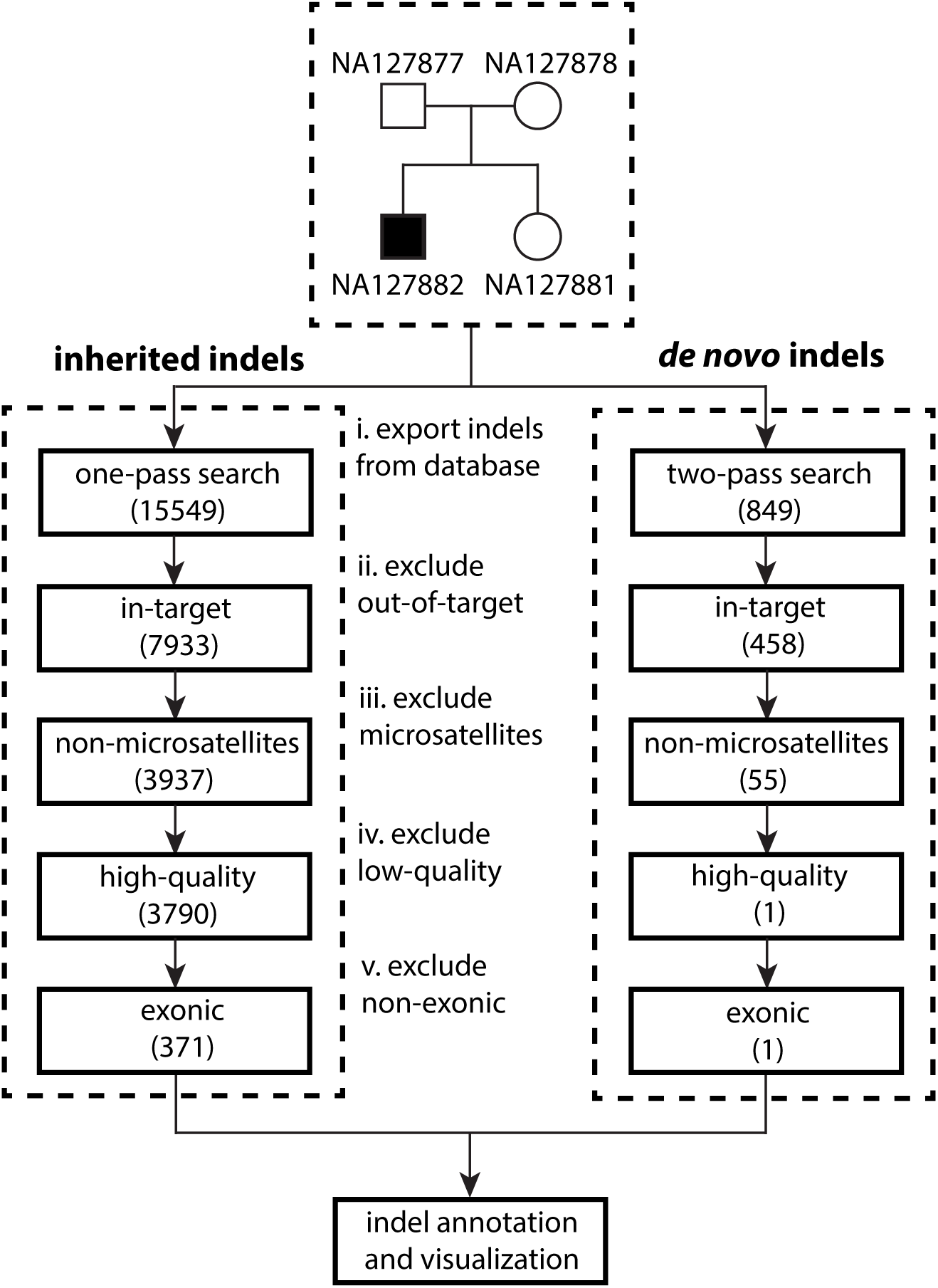
Filtering cascade of inherited and denovo indel calls. The numbers in each box denote the expected numbers of indel calls remained after filtering. The denovo indels were undergone a two-pass search to reduce the number of false positives.

#### Indel distribution within coding sequence

Because frameshift mutations can cause loss of function of a gene, these mutations are expected to be less frequent than frame-preserving mutations in the coding region. As shown in **Figure 11**, indels whose size is a multiple of three are much more abundant than others with similar sizes (+1 or −1).

**Figure 13.**
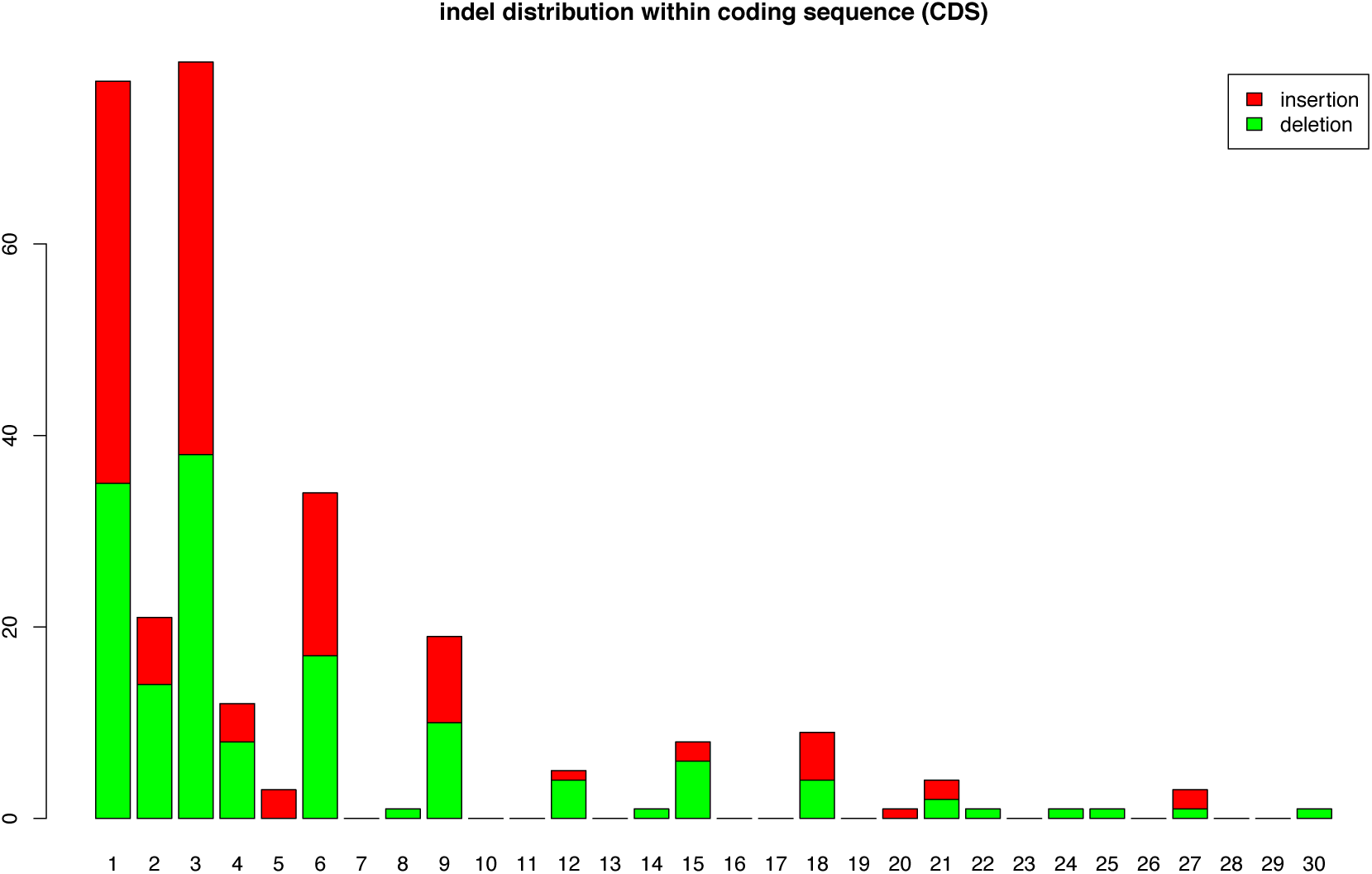
Frame-preserving indels are more abundant within coding sequences (CDS). Stacked histogram of insertions (red) and deletions (green) are shown in this figure.

#### A list of novel inherited frame-shift mutations in the family

Although this family has been investigated in many studies, many frameshift indels were not discovered in any public databases (**Table 3**). We observe a total of 6 novel frameshift mutations. Many of these indels are of a size larger than five bp. With the improvement of indel calling protocol introduced in this manuscript, we are able to identify these previously undiscovered loss-of-function mutations.

**Table 3.**
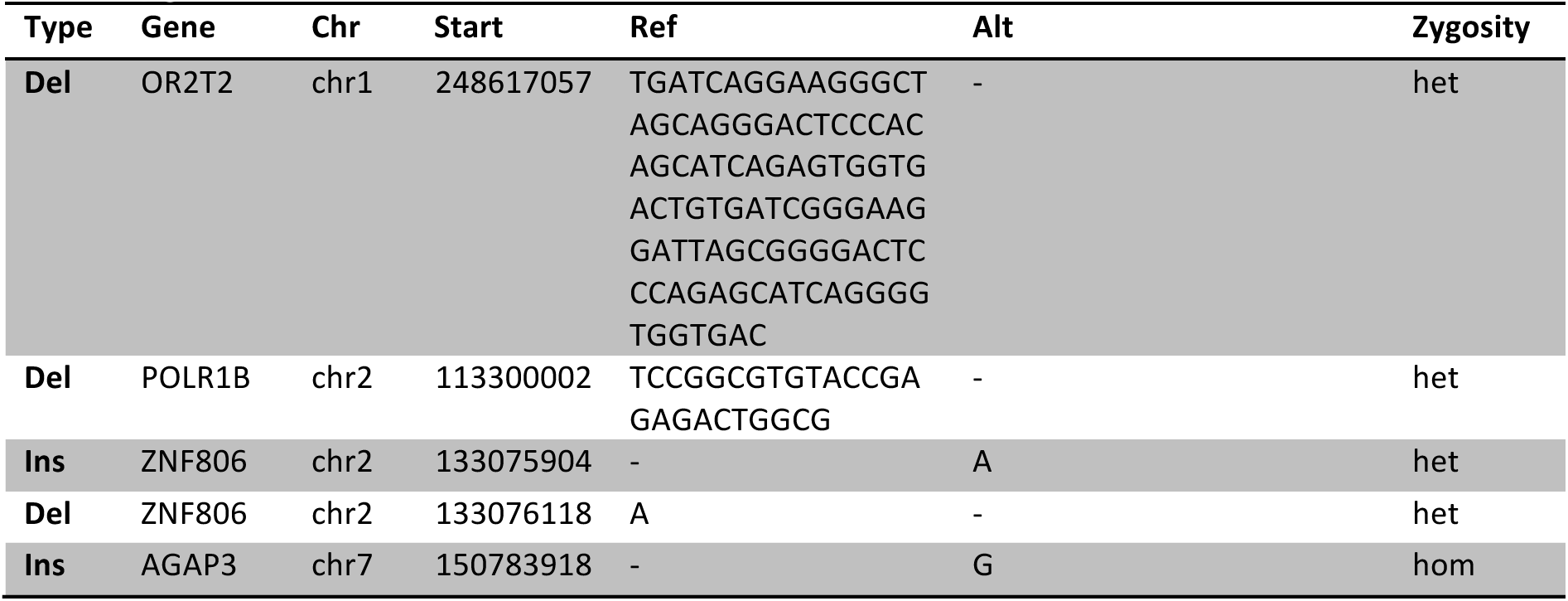
A selected list of inherited frameshift mutations that are not reported in 1000G ExAC, and ESP databases. Abbreviation: ins- insertion, del- deletion, hom – homozygous, het – heterozygous.

#### The de novo indel in the child and the alignment IGV screenshot

High-quality de novo indels usually share the following characteristics: 1) the number of reads in the region is close to the genome-wide mean coverage, 2) there are balanced number of reads supporting both the reference and alternative allele, 3) these indels are not located within or near short tandem repeat regions, 4) in the parents’ genome, there are no reads supporting the same indel presented in the child’s genome. **Figure 14** shows the screenshot of the IGV alignment of all four genomes. We can see a distinct signature of the deletion only presented in the affected kid, but not in anyone else in the family.

**Table 4.** Overview of the denovo deletion in the affected child

| Effect | Type | Gene annotation | Chr | Start | Ref | Alt | Zygoty |
| --- | --- | --- | --- | --- | --- | --- | --- |
| frameshift | deletion | HFM1:NM_001017975:exon4<br>:c.255delA:p.S85fs | chr1 | 9185<br>9889 | T | - | het |

**Figure 14.**
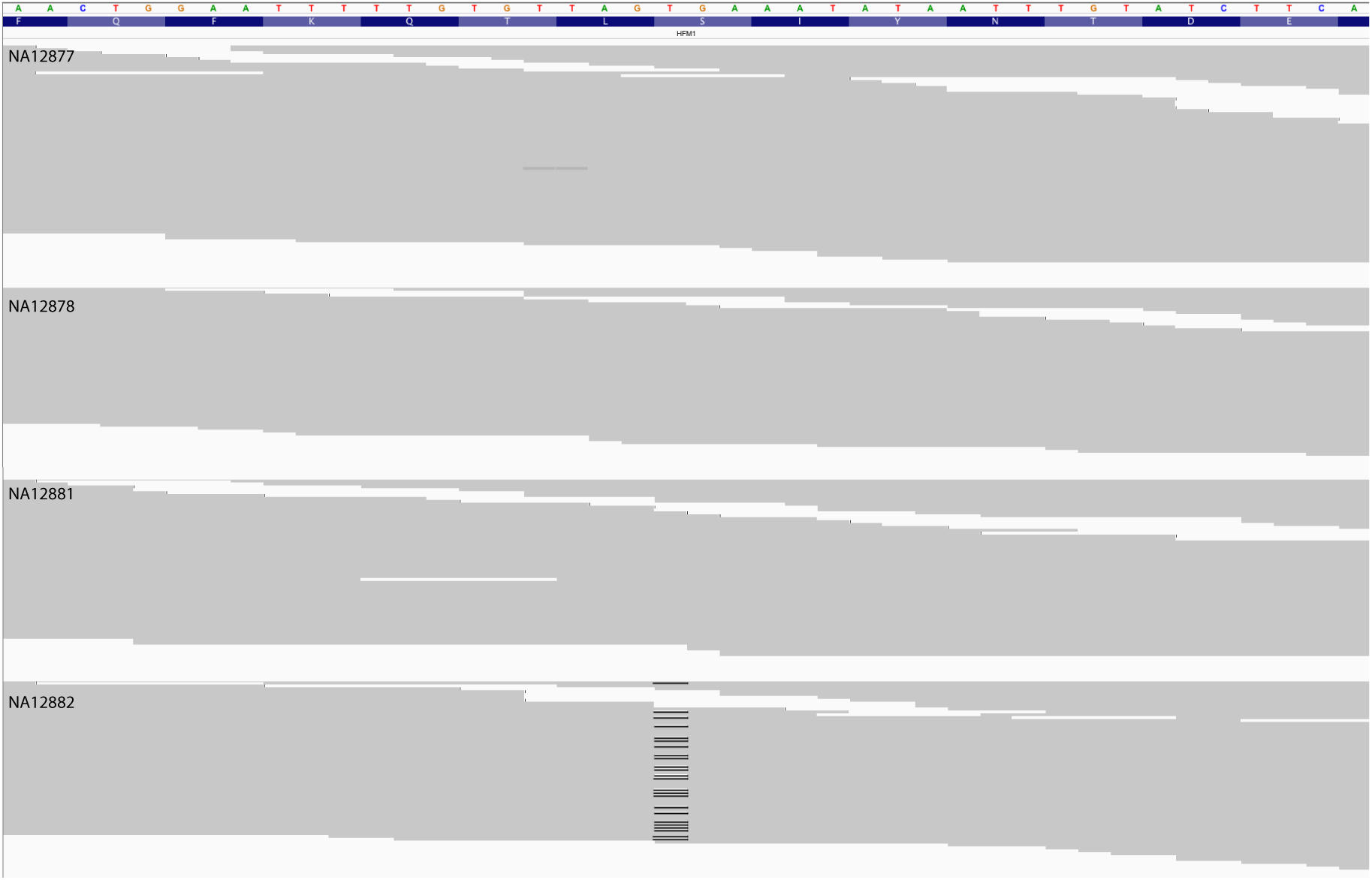
Screenshot of the alignment in IGV

## ACKNOLEDGEMENTS

The project was supported in part by the US National Institutes of Health (R01- HG006677) and US National Science Foundation (DBI-1350041) to M.C.S. and by the Cold Spring Harbor Laboratory (CSHL) Cancer Center Support Grant (5P30CA045508), the Stanley Institute for Cognitive Genomics and the Simons Foundation (SF51 and SF235988) to M.W.

## AUTHORS CONTRIBUTIONS

G.N. is the lead developer of Scalpel. M.C.S. contributed to the development of Scalpel and wrote the microsatellite detector scripts. H.F. contributed to enhance Scalpel, compiled the Scalpel resource bundle, and generated the figures in the manuscript. G.N., M.C.S., and M.W. conceived the Scalpel software project. G.N., M.C.S., and H.F. wrote the initial draft of the manuscript. M.C.S. is the Principal Investigator. All authors contributed to development and approved the final manuscript.

## Footnotes

1 http://vcftools.sourceforge.net/htslib.html#norm

2 https://www.broadinstitute.org/gatk/gatkdocs/org_broadinstitute_gatk_tools_walkers_indels_LeftAlignIndels.php

